# Direct observation of ATP-driven ubiquitin chain handling by Cdc48

**DOI:** 10.64898/2026.01.30.702746

**Authors:** D. Choudhary, J. R. Tait, M. Kracht, A. Roland, J. V. D. Boom, V. Sunderlíková, H. Meyer, S. J. Tans

## Abstract

Cdc48 (p97 or VCP in metazoans) targets polyubiquitin to selectively disassemble and degrade proteins. Cdc48 is believed to move along the ubiquitin chain towards linked proteins, but this has not been directly observed. By following single molecules in time, we find that the polypeptide branch points of ubiquitin chains are repeatedly inserted and rejected from the Cdc48 pore in ATP-driven manner, in bursts lasting up to seconds. This non-processive mode either ends by terminal substrate rejection, or advances to processive action, which drives ubiquitin unfolding in two steps, branch point translocation, and the extrusion of ubiquitin and linked protein as polypeptide loops in *trans*. Final retrograde movement can bring polypeptide segments back to *cis*. Our results establish the dynamics of ubiquitin chain handling by Cdc48, reveal key hallmarks of kinetic proofreading. We speculate that Cdc48 translocation may play a role in ubiquitin chain selection.

## Main Text

Cdc48 (p97 or VCP in metazoans) controls a striking array of cellular functions^1–4^. Notable examples are the degradation of misfolded proteins^2^, the remodelling of chromatin^5^ and membranes^6^, the regulation of DNA replication^7^, and the clearance of ribosomes^8^ and stress granules^9^. Cdc48 performs these tasks by disassembling protein complexes, extracting proteins from membranes, unfolding misfolded proteins, and by delivering proteins to the proteasome. A pivotal step is the selection of target proteins marked with polyubiquitin chains^10–13^. Despite the key role of structural changes of both Cdc48 and ubiquitin chains in this process, these dynamics have not been studied directly.

Cdc48 is believed to function by processively translocating polypeptides through its central pore^11,14–17^. This model is attractive, as it could explain how pulling forces are generated that unfold and extract polyubiquitinated proteins. However, the continuous, directional, ATP-driven movement that defines processive action has not been observed. Moreover, it is unclear whether processive translocation can explain how polypeptide branch points and loops are handled, which is necessary if Cdc48 moves along the polyubiquitin chain from its initial binding site to the linked protein^18^. Co-factors including Ufd1-Npl4, which both dock onto Cdc48 and bind ubiquitin moieties, are thought to control the selective targeting of ubiquitin chains^10,14^. Whether ATP-driven polyubiquitin handling steps after initial binding play a role is unknown.

Cryo-EM and bulk FRET studies have shown that initial binding to Ufd1-Npl4 causes one of the ubiquitin monomers to unfold and insert its N-terminus in the Cdc48 pore, without requiring ATP hydrolysis^10,14,19^, while leaving the first polypeptide branch point still on the *cis* side of Cdc48. The ATP-driven Cdc48 handling beyond this first interaction has been studied with biochemical approaches^4,12,18^, but has not been observed directly. Cryo-EM studies have further shown how Cdc48 contacts single linear polypeptides in its pore^14^, raising the question of how it handles the more complex polypeptide branch points and loops. Moreover, it was recently proposed that parts of the ubiquitin chain can move from *cis* to *trans* along the outside of Cdc48, using a transient opening of the Cdc48 ring^18^. Overall, the mechanisms of ubiquitin chain handling by Cdc48 are incompletely resolved. To address these questions, we developed a novel approach to directly measure Cdc48-induced conformational changes within individual ubiquitin chains in real time at the single-molecule level.

## Results

### Probing polyubiquitin conformation in real time

To detect Cdc48-generated conformational changes in polyubiquitinated substrates we used optical tweezers. For the latter, one typically attaches DNA handles to the N- and C-termini of the molecule of interest. However, due to its branched topology polyubiquitin has multiple termini which must remain free for insertion into Cdc48 (**Figure 1A**). To overcome these issues, we first used ubiquitin conjugation^20^ to grow a Lys48-linked ubiquitin chain on ubiquitin fused in-line to maltose binding protein (MBP; **Extended Data Fig. 1-3**). We terminated chain extension with a Lys48Arg ubiquitin mutant. Unique cysteines in this most distal ubiquitin and in MBP were used to attach two DNA handles (**Figure 1B**). The ubiquitin chain contains isopeptide bonds between all ubiquitin moieties, which we refer to as “branch points”. Single constructs were captured between two laser-controlled beads (**Figure 1C**). MBP was unfolded and kept that way at a low force of ∼10 pN, thus allowing the subsequent monitoring of the polyubiquitin chain length at nanometre resolution, and hence its conformational dynamics, in real time over extended periods (**Figure 1A**).

**Figure 1.**
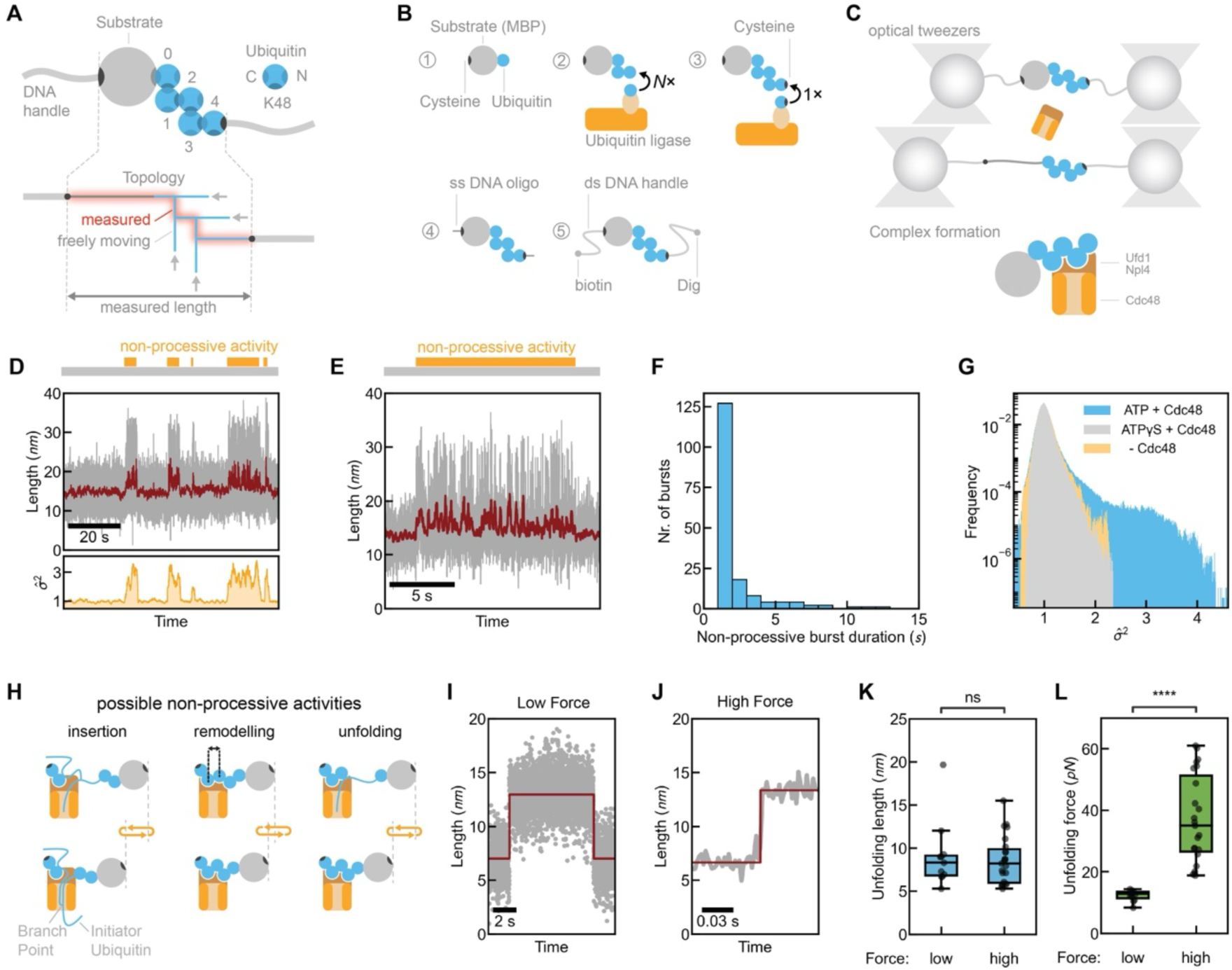
Non-processive Cdc48 activity in early handling stages. **A,** Tethered ubiquitin chain construct. DNA handle attachment points (black) allow end-to-end conformational probing of MBP and ubiquitin chain (red highlight). Ubiquitin C-termini are linked to Lys48 of subsequent ubiquitin. Arrows: N-termini free for Cdc48 pore insertion. The number of displayed ubiquitin moieties is chosen for illustrative purposes and does not reflect the studied chains (see Methods). **B,** Construction steps. (1) MBP-ubiquitin purification. (2) Ubiquitin ligation generating K48-linked chains. (3) Ligation of a distal ubiquitin K48R bearing an N-terminal cysteine (black) for handle attachment. (4) Linking of single-stranded DNA oligos. (5) Ligation of DNA handles. **C**, Measurement approach. Single constructs tethered between two laser-trapped beads are stretched to establish proper attachment, relaxed to low force (approximately 10 pN), and exposed to Cdc48, Ufd1, Npl4, and ATP regeneration system. Measured length informs on conformational changes induced by Cdc48. **D**, Top: construct length vs. time in presence of Cdc48 and ATP, showing non-processive Cdc48 activity. Top: Raw (grey) and filtered (red, Methods) length signal. Bottom: σĮ^2^ that quantifies the fluctuation intensity (Methods) vs. time, which identifies the non-processive bursts. **E**, Zoom of non-processive activity. **F**, Burst duration histogram. The near-exponential distribution is consistent with single Cdc48 molecules stochastically switching to a non-active state. N_bursts_ = 173, N_molecules_ = 62, of which 34 % contain a burst. **G**, Distribution of σĮ^2^ that quantifies the fluctuation intensity (panel D, bottom; Methods). Activity bursts with Cdc48 and ATP lead to values of σĮ^2^ > 1 and a skewed distribution (skewness 10.8), not observed for Cdc48 with ATPγS, or without Cdc48 (skewness 1.3 and 1.4 respectively). **H,** Possible length change mechanisms. The initiator ubiquitin is visualized at an arbitrary position, and could instead be more distal along the ubiquitin chain. **I,** Example trace of a single ubiquitin (un)folding at 10 pN and constant trap position. **J**, Example trace of ubiquitin unfolding during force ramp force. **K**, Corresponding measured length changes. Number of unfolding events, N_events_ = 11 and 17 respectively. N_molecules_ = 8 and 23 respectively. **L**, Corresponding unfolding forces.

### Non-processive Cdc48 activity in early handling stages

Introduction of Cdc48 with Ufd1-Npl4 and ATP led to bursts of length fluctuations that lasted up to seconds (**Figure 1D-E**, **Extended Data Fig. 4E-F**). Bursts were identified automatically by analysing the fluctuation intensity (**Figure 1D**, bottom; Supplementary Information). Several features suggested the bursts were generated by single Cdc48-Ufd1-Npl4 complexes (hereafter referred to as Cdc48): the bursts started and stopped in a discrete manner (**Figure 1D**, top); they showed constant fluctuation intensity, both during one burst and between subsequent bursts (**Figure 1D**, bottom); and their durations were nearly exponentially distributed (**Figure 1F**), as expected for single molecules crossing an activation barrier^21^. The bursts were observed for 34% of the tethered substrates, after a waiting period that was broadly distributed (**Extended Data Fig. 4A**). The other tethered substrates did not show Cdc48-mediated activity. However, once a burst was detected, the chance of a subsequent burst was high (95%). More specifically, the characteristic waiting time for subsequent bursts (about 3 s, **Extended Data Fig. 4B, G**) was 58 times shorter than the waiting time for the first burst (about 175 s., **Extended Data Fig. 4A, G**). A parsimonious explanation is that one Cdc48 hexamer drives multiple bursts. However, one cannot formally exclude that Cdc48 unbinds and another Cdc48 hexamer binds. Note that this scenario would require ubiquitin-cofactor dissociation and initiator ubiquitin release from the Cdc48 pore, and yet unknown mechanisms for subsequent engagements being more frequent than first engagements. Overall, these data indicated the formation of Cdc48-polyubiquitin complexes that are activity-competent, followed by stochastic switching between active and non-active states.

The length fluctuation were 7.6 ± 6.0 nm in magnitude (mean and SD, **Figure 1D-E**, **Extended Data Fig. 4C**). We wondered if they could reflect remodelling of the polyubiquitin structure by Cdc48 (**Figure 1H**, middle), given that di-ubiquitin can alternate between closed and open states^22–24^. However, this switch yields a length change of about 0.77 nm only for di-ubiquitin, or 1.5 nm for tri-ubiquitin (**Extended Data Fig. 5**). The latter is effectively the maximum because only a few ubiquitins interact with Ufd1-Npl4-Cdc48, and the fluctuations depended strictly on Cdc48. Ubiquitin unfolding can yield length increases of about 8 nm (**Figure 1I-L**, **1H**, right). However, ubiquitin is known to be highly force stable^25,26^, and indeed typically only unfolded when pulled to well over 20 pN in our assay (**Figure 1J-L**). At the 10 pN tension, ubiquitin unfolding was very rare (0.01 events per minute on average), and rather occurred as single discrete transitions between distinct states as usual for (un)folding events^27^ (**Figure 1I**).

Activity bursts were not observed when ATP was exchanged for the non-hydrolysable ATPγS (**Figure 1G**), indicating a link to translocation. Insertion and release of the first branch point would indeed lead to length decreases and increases (**Figure 1H**, left). To further test ATP dependence, we studied a Walker B mutation (E315Q) that disrupts ATP hydrolysis of the D1 ring but not ATP binding, and still facilitates N-terminal insertion of the initiator ubiquitin^10,14,19,28^. We found that the non-processive fluctuations were abrogated for this mutant, as well as for WT in the APO state (**Extended Data Fig. 4D**). While other mechanisms cannot be formally excluded from these data, they suggest that active translocation drives the non-processive fluctuations, as well as a role for the D1 ring. Note that this translocation activity could in turn cause secondary conformational changes in ubiquitin, including its (partial) folding and unfolding, which may also contribute.

### Cdc48 switches to processive translocation

We surmised that the polyubiquitin conformation could play a role in the lack of processive Cdc48 activity, as spontaneous conformational changes of di-ubiquitin have for instance been shown to limit deubiquitinase activity^23,24^. Testing this idea is non-trivial, as these nanoscale conformational dynamics are too small to measure with optical tweezers. Moreover, different conformations of otherwise identical ubiquitin chains and their dynamics are poorly understood and, cannot be controlled experimentally by generating different types of chains. Attempting to limit the polyubiquitin conformational changes by lowering the applied force also does not work, as the resulting large increases in measurement noise render our detection of activity impossible. To overcome these issues, we altered the DNA handle sites such that the ubiquitin chain was conformationally free (**Figure 2A-B**, **Extended Data Fig. 1-3, 6-8**). Note that the ubiquitin chain is no longer probed at its end, and hence not all its length changes are detected. However, the construct was designed such that the most proximal ubiquitin (Ubi_0_) and MBP are still probed end-to-end and hence their length changes are still measured, which is central here. Two ubiquitin chains were incorporated to optimize the assay (**Figure 2A-B**).

**Figure 2.**
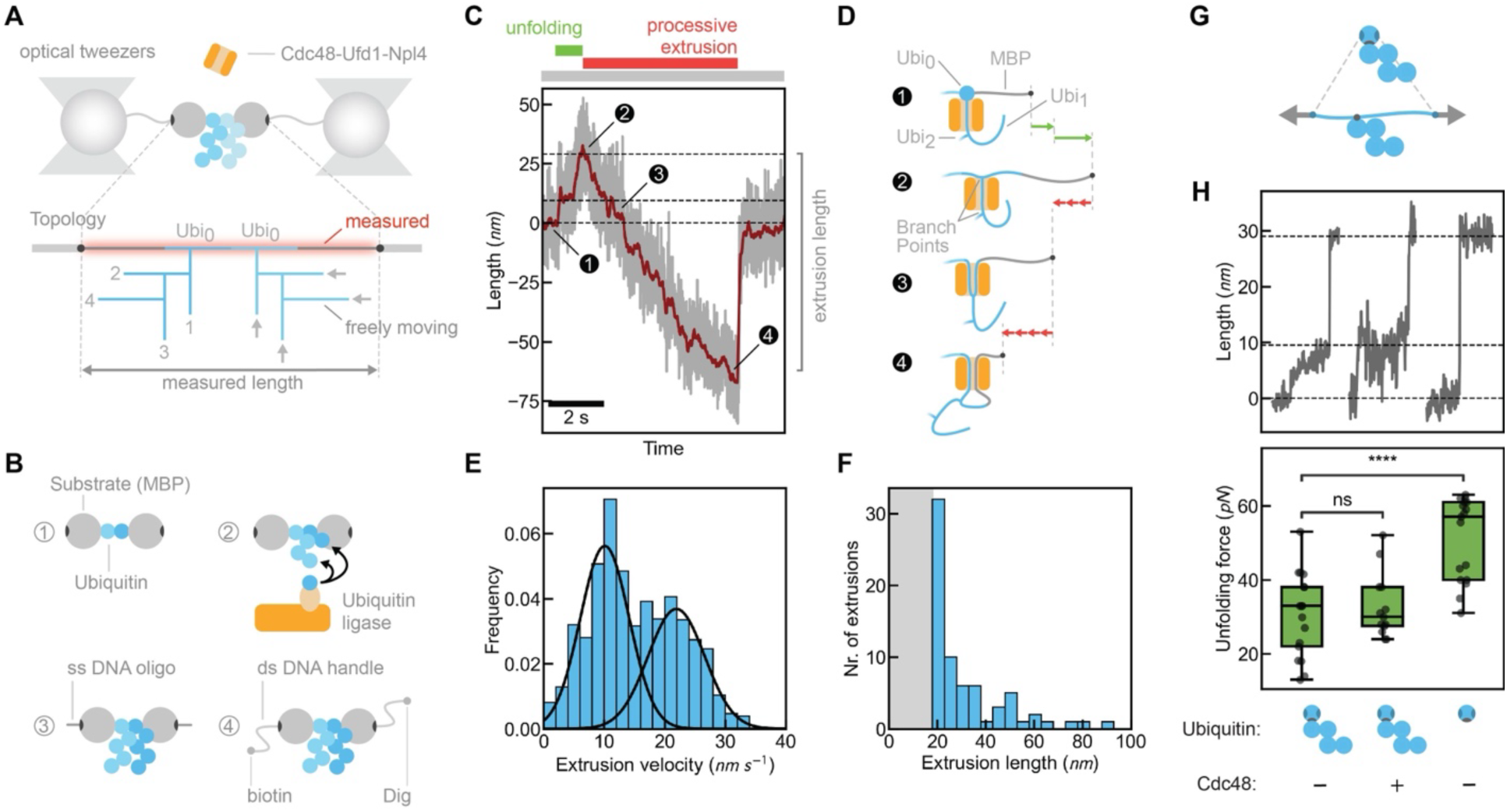
Processive extrusion of polyubiquitin and substrate loops by Cdc48. **A,** Optical tweezers assay for conformationally-free construct. DNA handles are attached such that MBP and Ubi_0_ are probed (red highlight) while the rest of the ubiquitin chain is free to move. Two MBP-ubiquitin chain copies are included to optimize the assay. Numbers indicate ubiquitin moieties, arrows are N-termini free for Cdc48 pore insertion. **B**, Construction steps. **C,** Construct length vs. time with Cdc48 and ATP, showing processive Cdc48 activity. Numbers: see panel D. Dashed lines: ubiquitin (un)folded states. Raw and filtered length signals are shown in grey and red respectively. **D,** Illustration corresponding to panel c. **E,** Distribution of extrusion velocity. Black lines: Gaussian Mixture fit showing peaks at approximately 10 and 23 nm/s, consistent with one-and two-arm translocation. **F,** Distribution of extrusion lengths, determined by automated analysis. Grey: detection limit. Lengths above 50 nm show substrate extrusion, even if both Ubi_0_ moieties would be fully translocated first. N_events_ = 64, N_molecules_ = 60, of which 27 % contain an extrusion. **G,** Cartoon of ubiquitin unfolding experiments. **H,** Unfolding of tethered ubiquitin during force ramp. Left: with ubiquitin chain, without Cdc48. Middle: with ubiquitin chain, with Cdc48. Right: without ubiquitin chain, without Cdc48. Top: example unfolding traces. Bottom: Force of ubiquitin unfolding. Number of unfolding events (N_events_) is: 21, 12, and 16 respectively. Number of probed ubiquitin moieties (N_molecules_) is: 193, 94, and 204 respectively. Fraction of unfolded ubiquitin moieties is: 11%, 13%, and 8% respectively. Unfolding is shown to occur in two-step manner and lower-force unfolding when ubiquitin chain is added.

Exposure to Cdc48 now yielded continuous decreases in length (**Figure 2C**). The length changes were unidirectional (**Figure 2C**), showed roughly constant slopes (**Extended Data Fig. 9**), and reached lengths of up to 100 nm (**Figure 2F**). These data also indicated the extrusion of polypeptide loops by Cdc48 (**Figure 2D**): with both substrate termini tethered, the length decreases must involve the formation of a polypeptide loop. Given a 2-residue translocation step size per Cdc48 monomer^29^, Cdc48 thus catalysed multiple steps in sequence over multiple cycles of ATP hydrolysis without releasing the substrate, which is the hallmark of processive action. Indeed, Cdc48 must maintain its grip on the polypeptide loop in order to drive the length decreases, as releasing grip would allow the loop to extend again. Polypeptide loop extrusion was further supported by the presence of two peaks, at velocities *v* (10.0 nm/s) and 2*v* (21.7 nm/s) (**Figure 2E**), which is consistent with one or two arms (polypeptide segments) of the loop being driven through the pore respectively^30^. The experiments also demonstrated that Cdc48 performs mechanical work, because they show driven movement in a direction opposite to the external force. This capability is relevant to the unfolding of ubiquitin that we investigated next.

### Cdc48 processivity drives two-step ubiquitin unfolding

Our model (**Figure 2D**) suggested that Cdc48 must first unfold Ubi_0_ to be able to translocate Ubi_0_ as a loop. We indeed observed discrete increases in length before the gradual decreases (**Figure 2C**, point 1 to 2). These increases totalled 28.4 ± 0.5 nm, in line with the estimated ubiquitin length (26 nm). However, they occurred in two steps, which is inconsistent with the main one-step ubiquitin unfolding pathway previously reported^27^, while two-step unfolding was observed in a minority (5%) of cases^31,32^. Next, we studied the unfolding of ubiquitin without an attached polyubiquitin chain. Ramping the force in absence of Cdc48 indeed yielded abrupt length increases of 28 nm in a single step (**Figure 2H**, right). After ligating a (freely moving) ubiquitin chain, the 28 nm transition occurred in two steps of about 10 and 18 nm, as observed with Cdc48 (**Figure 2C, H**, top). The average measured unfolding force also decreased: from 45 pN for single ubiquitin to 32 pN when polyubiquitin was ligated (**Figure 2H**, bottom). Note that most ubiquitin moieties did not unfold below the maximum force of 65 pN that we can apply due to DNA melting. Ubiquitin unfolding forces above 200 pN have been reported^25,27^, though these studies used AFM which use higher loading rates and typically measure higher unfolding forces. This finding indicated a decreased stability^33^, as well as an altered unfolding pathway of ubiquitin moieties, when they reside within ubiquitin chains. Further, the data corroborated that the two-step length increase seen prior to extrusion (**Figure 2C**) reflected Cdc48-driven ubiquitin unfolding.

### Ubiquitin branch points and loops are translocated processively

When Cdc48 has just unfolded Ubi_0_, a ubiquitin branch point is expected to be situated at the *cis* entry of the Cdc48 pore (**Figure 2D**, point 2). Once the measured length has decreased by twice the pore length, about 18 nm^14,34^, the branch point has moved to the *trans* side of Cdc48 (**Figure 2C**, point 3; **Extended Data Fig. 10, 15**). Note that the more distal ubiquitin monomers are either still bound to Ufd1-Npl4 in *cis*, or moved to *trans* via the proposed Cdc48 ring opening^18^. The average velocity during branch point traversal was 31.5 nm/s, at the high end of the velocity distribution (**Figure 2E**). Once successfully inserted for processive translocation, the branch point – with its bent Ubi_0_ polypeptide segment – thus did not cause a major reduction in translocation velocity. Further translocation caused polypeptide loop extrusion in *trans*. The extrusion can continue beyond Ubi_0_ to include MBP, as extruded lengths were observed that exceeded the ubiquitin lengths, even as many extruded lengths were less than 40 nm (**Figure 2C, F**).

The extrusion length is roughly exponential (**Figure 2F**), consistent with random events in the complex triggering the end of extrusion (see below). The measured extruded length can also be limited by our applied forces, though we note that counteracting forces can also occur *in vivo*, for instance when Cdc48 pulls on folded structures it encounters. Further decreasing the applied force leads to large measurement noise that precludes extrusion detection. Methods such as FRET may overcome this issue, but bring other unresolved challenges. The extruded length could also be limited by an obstacle, which here may be the second ubiquitin chain that Cdc48 encounters as it translocates Ubi_0_. To test this idea, we generated a construct with only one Ubi_0_ copy, and thus one ubiquitin chain (**Extended Data Fig. 11A**). We again found processive movement (**Extended Data Fig. 11B-D**), and the observed extruded lengths were consistent with the previous data (**Figure 2F**). The extrusion velocity histogram again showed two peaks, one having twice the velocity of the other (**Extended Data Fig. 11C**). Hence, these data indicated that the second Ubi_0_ branch point is not a main trigger of translocation arrest. Overall, our data showed that processive translocation drives the handling of polyubiquitin loops and branch points by Cdc48.

### Retrograde movement through the Cdc48 pore

Extrusions typically ended how they started: with sudden length increases (**Figure 2C**, **3A**). The length changes correlated with the previous extrusion-driven length decreases (**Figure 3B**). In principle, opening of the Cdc48 ring^18^ could extend the extruded polypeptide loops, and hence yield such correlations. Such a ring opening and lateral peptide release would break the loop and render subsequent extrusions by the same Cdc48 topologically difficult. However, we did typically observe a translocation event after the length increase (91% of events; **Figure 3A)**. The data were in line with Cdc48 pore loops releasing their grip on the substrate, the extruded polypeptide segment moving backwards through the Cdc48 pore, and subsequently translocation restarting (**Figure 3A, D**, cartoons). Note that the applied force here acts continuously, and can draw all the translocated Ubi_0_ polypeptide back to cis, which is likely different *in vivo* where Cdc48-generated and entropic forces can exist but not as continuously. Also note that in our experiments, only Ubi_0_ should be drawn back to *cis*, with the Ubi_1_-Ubi_0_ branch point again positioned at the Cdc48 pore entry (**Figure 2D**, point 2). The characteristic waiting time between subsequent (extrusion) events (about 9 s, **Extended Data Fig. 12**) was again an order of magnitude smaller than for the first event (about 208 s). Hence, these data indicated the following picture: once a Cdc48 hexamer completed the (comparatively long) process of binding the ubiquitin chain, reaching and extruding Ubi_0_, and releasing (part of) Ubi_0_ again to *cis*, Cdc48 can initiate subsequent extrusions faster.

**Figure 3.**
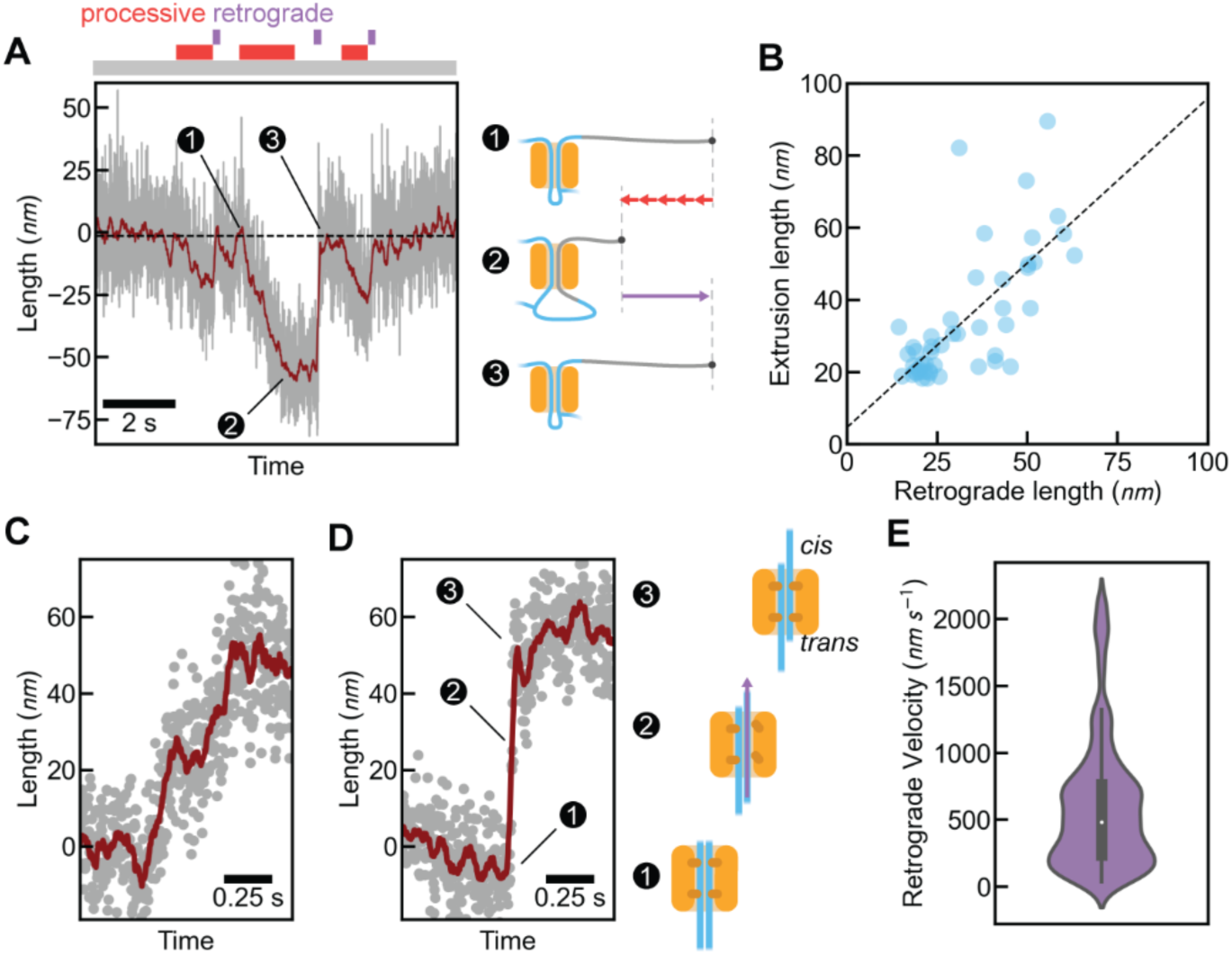
Retrograde movement through the Cdc48 pore. **A,** Length vs. time for conformationally-free construct (Figure 2A-B) in presence of Cdc48 and ATP, showing retrograde movement of translocated polypeptide segments. Retrograde movement is often followed by a subsequent extrusion event. **B,** Extruded length vs. length of subsequent retrograde movement. Linear fit has coefficient of determination *R*^2^= 0.51. **C,** Example of a slow retrograde movement. **D,** Example of a fast retrograde movement. **E,** Observed retrograde velocities. N_events_ = 79, N_molecules_ = 62, of which 41 % contain a retrograde movement. Across panels A-C, the raw and filtered signals are shown in grey and red respectively.

Automated analysis revealed the retrograde velocity was broadly distributed (**Figure 3E**) and included slow as well as fast length increases (**Figure 3C-D**). These data suggested friction with the inner Cdc48 surface as the polypeptides moved back to *cis*, possibly involving interactions with the Cdc48 pore loops. In our model, the polypeptide loop on the *trans* side of Cdc48 contains a polypeptide branch point (**Figure 2D**, **3A**). Consistently, we detected transient arrests during the retrograde movement when this branch point was expected to encounter the *trans* exit of the Cdc48 pore (**Extended Data Fig. 10**, point 4). Overall, the data showed that Cdc48 can transiently release all contacts to one of the peptide segments, allowing its substrates to move back to *cis*, while Cdc48 remains engaged with the substrate for further processing.

### Non-processive Cdc48 activity in late handling stages

We showed that Cdc48 acts non-processively in early polyubiquitin handling stages (**Figure 1**). Similar behaviour was observed in later stages: bursts of length fluctuations were observed when a (proximal) branch point was situated at the *cis* entry of Cdc48, and the corresponding ubiquitin moiety in *cis* was unfolded (**Figure 4A**). As during the early stages (**Figure 1**), the bursts could start and end in discrete manner and last for seconds (**Figure 4A**) and were not observed in absence of Cdc48 (**Figure 4B**). These data showed that the repeated insert-reject cycles and processive extrusion followed each other directly (**Figure 4A**). Moreover, after retrograde movement we found Cdc48 molecules that were still actively handling the substrate, in both the processive and non-processive modes (**Figure 3A**, **Figure 4A**). This is consistent with our model: retrograde movement brings Cdc48 back to the same general position on its substrate where processive activity was initiated, namely with the branch point at the *cis*-side of the Cdc48 pore.

**Figure 4.**
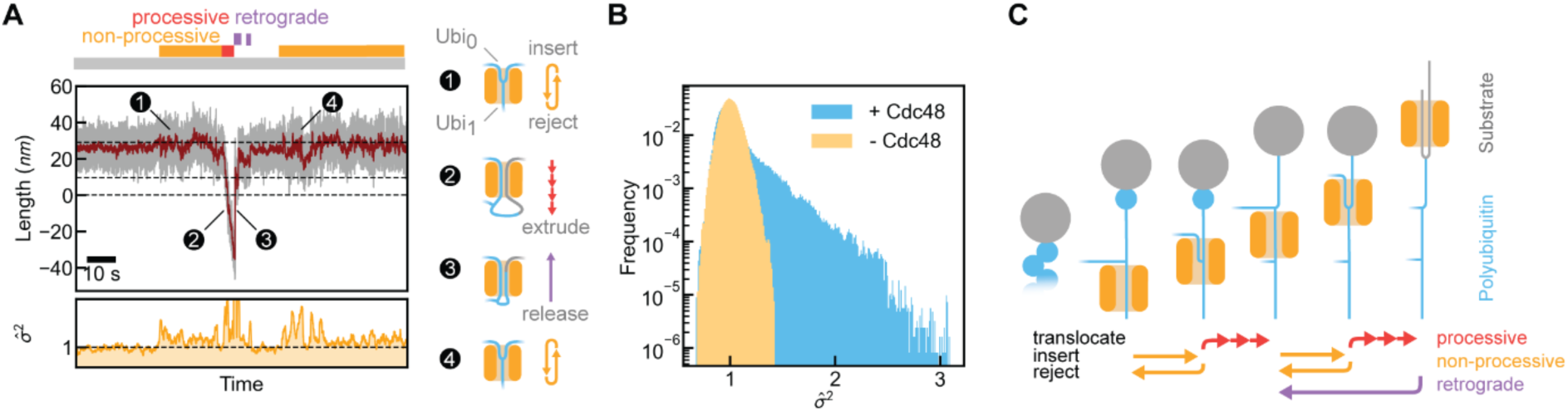
Non-processive Cdc48 activity in late handling stages. **A,** Top: length vs. time for conformationally-free construct (Figure 2A-B) in presence of Cdc48 and ATP. (1) non-processive Cdc48 activity, (2) processive extrusion, (3) retrograde movement, (4) non-processive activity. Dashed lines indicate ubiquitin unfolded states. Raw and filtered signals are shown in grey and red respectively. Bottom: σĮ^2^ that quantifies the fluctuation intensity (Methods) vs. time, showing fluctuations increase before processive translocation. **B,** Distribution of σĮ^2^ (Methods). Without Cdc48, the distribution is nearly Gaussian (skewness 0.3). Non-processive Cdc48 bursts cause a shoulder at higher enrichment (skewness 2.8). **C,** Schematic overview of observed polyubiquitin handling by Cdc48 with key hallmarks of kinetic proofreading. Left: when binding the initiator ubiquitin, the latter unfolds and N-terminally inserts in Cdc48 pore in ATP-independent manner^1–3^. Cdc48 encounters polypeptide branch points when moving along the ubiquitin chain toward target protein (grey). Repeated rejection of branch points from the Cdc48 pore after insertion (yellow arrows) either ends with terminal Cdc48 arrest, or is followed by processive translocation (red arrows), which drives ubiquitin unfolding and loop extrusion. Such repeated rejection conditionally followed by reaction progression is the central hallmark of kinetic proofreading. The initiator ubiquitin is visualized at an arbitrary position, and could instead be more distal along the ubiquitin chain.

## Discussion

By measuring conformational dynamics in time, we directly followed the ATP-driven handling of ubiquitin chains by Cdc48 (**Figure 4C**), which occurs after the ATP-independent initial binding and N-terminal insertion of one of the ubiquitin moieties, as shown previously by Cryo-EM and population-level FRET methods^10,14,19^. Our data revealed processive and non-processive modes that together underlie this ATP-driven polyubiquitin handling. The directional, continuous, constant-velocity length changes unambiguously showed processive Cdc48 function. Processive translocation of one ubiquitin moiety was found to generate forces that unfold the next ubiquitin. Ubiquitin unfolded *via* a two-step pathway and at lower force, which were induced by the ubiquitin chain itself and indicated conformational remodelling in the chain. Ubiquitin unfolding yielded a polypeptide branch point at the *cis* entry of the Cdc48 pore. Notably, the polypeptide branch points of ubiquitin chains were repeatedly rejected from the Cdc48 pore after insertion. This non-processive activity of Cdc48 occurred in bursts lasting up to tens of seconds, which ended either in terminal Cdc48 rejection or in advancing to translocation. In the latter case, Cdc48 action was highly processive, driving the rapid traversal of branch-points to the pore exit, and the extrusion of ubiquitin and substrates as polypeptide loops in *trans* (**Figure 4C**).

ATP-driven trial-reject cycles with conditional reaction progression are features observed in kinetic proofreading. The observed mechanism also shows aspects that differ from proofreading systems like ribosomes, which sample and reject tRNAs to ligate amino acids with high fidelity^35,36^. Cdc48 rather rejects polypeptide branch points, and may encounter multiple branch points along the ubiquitin chain^37,38^. The data implied that conformational changes in the ubiquitin chain affected rejection, as the conformationally free chains advanced to processive extrusion (**Figure 2, 3**, and **4, Extended Data Figure 11**) while the tethered ones did not (**Figure 1**). In similar vein, deubiquitinase activity was shown to depend on di-ubiquitin conformational states, which are populated stochastically^39,40^. Unlike this conformational selection, Cdc48 showed the energy-driven repetitive cycles that typify kinetic proofreading. Diverse parts of the Cdc48-substrate complex could play a role here: the conformations of the ubiquitin moieties distal to the initiator ubiquitin or of the polypeptides that form the branch point, and how these parts contact the Cdc48 pore loops, the Ufd1-Npl4 cofactors, or other regulators.

In the current paradigm, initial binding by Cdc48/p97 cofactors controls ubiquitin chain selection. Yet, while the known cofactors show specificity in binding different ubiquitin chains, they may not fully explain how the wide range of ubiquitin chains and their divergent fates (proteasome, Cdc48, de-ubiquitination) can be controlled^13,41^. Our data opens up the possibility that ubiquitin chain translocation by Cdc48/p97 may not serve only to move from the initial binding site to the target protein, but also to affect substrate selection. One may further speculate that Cdc48/p97 leverages the ubiquitin conformational dynamics, and the related decreases in ubiquitin unfolding forces^23,24,33^ observed here. Regulatory cofactors and ubiquitin modifications could for instance affect selection by influencing polyubiquitin conformation. As such, our findings could be relevant to the handling and selection of ubiquitin chains with heterotypic linkages^13^ or SUMOylation and phosphorylation modifications^42^, ubiquitination and de-ubiquitination balance^43^, and proteasome and Cdc48/p97/VCP competition^44^.

Our data further showed translocated polypeptide segments that were released back to the *cis* side of Cdc48. The pore loops that hold polypeptides within the Cdc48 pore can thus dissociate simultaneously. Whether this involves a coordinated conformational change of Cdc48 monomers is unclear. Transient rebinding of the pore loops, and resulting effective friction, may explain the observed slow retrograde movements. The functional role of this retrograde movement is an open question. It is consistent with the rejection step of the non-processive Cdc48 action, may allow Cdc48 to disassemble protein complexes without the need to fully translocate substrates^12,44^, and offer translocation and unfolding retries for poorly inserted or difficult to unfold substrates.

The *cis* release measured here concerns released polypeptide segments, and do not correspond to a complete dissociation of ubiquitinated substrate from Cdc48. Indeed, retrograde movement is often followed by translocation events, indicating the substrate is not fully dissociated. Complete release may be suppressed by ubiquitins in *trans*, which can arise by *trans* refolding, ubiquitin chain elongases acting in *trans* such as Ufd2^45^, and by proposed Cdc48 ring opening^18^. Conversely, complete substrate release may be promoted by deubiquitinases including Otu1^23,24,45,46^, and ubiquitin binding proteins like Rad23 that can direct proteins to the proteasome^47,48^. Retrograde movement can be promoted by our applied forces, for instance by increasing the pore-loop dissociation rate. At the same time, pN level forces can be generated *in vivo* by Cdc48 pulling on folds that it encounters, as well as by entropic pulling^49^ and *cis* refolding^50^, which may thus also promote pore loop dissociation. Here we also show directly that Cdc48 translocation can unfold proteins such as ubiquitin, which also leads to opposing forces, and suggests they do not readily trigger pore loop dissociation. It is possible that other factors further enhance processivity *in vivo*.

The backward *cis* release and forward translocation of branch points and loops also raises several structural questions, as Cryo-EM studies have only shown single polypeptides in the Cdc48 pore^14–17^. It is also not fully clear what sets the scale of the non-processive movements, in which the ATP-driven movements of the D1 pore loops play a key role. Their maximum amplitude appeared rather constant in time within and between bursts, while amplitude variations were observed. The on average 7.6 nm length changes imply branch point movements of half that, which would roughly match the distance between the pore entry and the D1 pore loops, or the distance between the D1 and D2 pore loops. Hence, branch point rejection may be triggered by a failure to pass the D1 or D2 pore loops, though we stress that these are speculations and other potential triggers like ubiquitin moieties moving into contact with Cdc48 components could play a role. Furthermore, it is unknown how Cdc48 can switch between holding and moving one or two polypeptide segments, as shown in our data. Finally, it remains an open question whether the repetitive pulling motions during non-processive action allow protein complex disassembly with minimal substrate unfolding.

Our data highlight the critical role of temporal dynamics in ubiquitin chain handling by Cdc48. The observed mechanisms may affect Cdc48 functions that are regulated by other cofactors, like membrane fusion mediated by p47/Shp1^51,52^, DNA repair mediated by DVC1 and Wss1^1,5^, autophagy mediated by UBXD1^9,53^, chromatin remodelling mediated by UBXN7/Ubx5^1^, and ribosome quality control mediated by Vms1 and Rqc1^8,54^.

## Acknowledgements

We thank Pieter-Rein ten Wolde, Tom Shimizu, Kristina Ganzinger, and Marija Iljina for comments and critical reading of the manuscript; and all members of the Tans lab for helpful discussion.

## Author contributions

DC, AR, HM and SJT conceived and designed the research; AR, VS, HM and SJT designed and generated the substrate constructs; DC and AR performed the optical tweezers experiments; MK, JVDB and VS purified and generated the substrate and Cdc48 proteins; JRT wrote the data analysis software; DC, JRT, AR, and SJT analysed the data; and DC, JRT, MK, JVDB, HM, and SJT wrote the manuscript.

## Data and materials availability

The data that support the findings of this study are available within the paper and the Supplementary Information files.

## Code availability

Code for data analysis (Cdc48 activity) is available at https://github.com/tait-py/KineticProofreadingCdc48.

## Methods

### Protein expression and Purification

#### Expression and purification of substrates

A terminal cysteine residue was introduced at the free end of MBP in the expressed MBP-ubiquitin (hereafter MBP-ubi) and MBP-ubiquitin-ubiquitin-MBP (hereafter MBP-ubi-ubi-MBP) using the pET28 vector. The two constructs were expressed in *Escherichia coli* BL21(DE3) cells (New England Biolabs) in LB medium containing kanamycin (50 μg/ml) overnight at 18°C. Cell pallets were collected through centrifugation at 5000g for 20 minutes at 4°C. Subsequently, the pellet was resuspended in ice-cold buffer (50 mM phosphate buffer pH 7.5, 200 mM NaCl, 10 mM EDTA, 50 mM glutamic acid–arginine and 1 mM DTT) then lysed via Emulsiflex homogenizer. Clear lysate was procured by 60 min centrifugation at 50,000g at 4°C. The protein was allowed to bind to amylose resin (New England Biolabs) for 1 hour at 4°C with slight agitation. In order to remove unspecific bound proteins and nucleic acid, the column was washed 4 times in cold washing buffer (50mM Tris-HCl pH 7.5, 200 mM NaCl, 10 mM EDTA and 1mM DTT). The protein was eluted with 20 mM maltose. The resin was allowed to incubate in buffer NAME for 20 minutes at 4°C prior to this stage. The protein was thoroughly washed with Tris buffer and supplemented with 20 mM maltose.

#### Plasmids

A hexa-histidine tag and a cysteine were added in front of the N-terminus of ubiquitin in pET23a Ubiquitin K48R by mutagenesis PCR. Plasmid pET28 His-gp78-ubc8^20^ was received as a gift from Raymond Deshaies’ lab. Plasmid K27Sumo His-SUMO-Cdc48^55^ was received as a gift from Tom Rapoport’s lab. Plasmid pET28 His-mUbe1^56^ was purchased from Addgene (plasmid # 32534). Yeast ORF UFD1 (YGR048W) and yeast ORF Npl4 (YBR170C) were purchased from GE-Dharmacon and cloned into pET41, with a C-terminal His-tag for Ufd1, using NdeI and XhoI (Ufd1) or NdeI and EcoRI (Npl4) respectively. The construct pET28 His-ubi-ubi-mEos3.2 was adapted from Blythe et al, by replacing GFP with mEos3.2^20^. Human YOD1 and Ataxin-3 were cloned into pET28 with an N-terminal His-tag.

#### Ubiquitin chains and ubiquitin antibodies

Distinct ubiquitin chains (K48 3-7 UC-220, K63 3-7 UC-320) were purchased from Boston Biochem. Chain type specific ubiquitination was detected using the following antibodies from Merck Millipore: α-pan-ubiquitin FK2 mouse monoclonal (04-263), α-K48-ubiquitin Apu2 rabbit monoclonal (05-1307), α-K63-ubiquitin Apu3 rabbit monoclonal (05-1308).

#### Expression and purification of Cdc48 and co-factors

Proteins were expressed in Bl21(DE3) cells. Cultures were grown at 37 °C until an OD600 of 0.6, induced with 0.5 mM ITPG and incubated overnight at 18 °C. Cells were harvested, resuspended in lysis-buffer (50 mM HEPES pH 8.0, 150 mM KCl, 2 mM MgCl_2_, 5% glycerol) and stored at - 80°C.

Proteins carrying a poly-Histidine tag were purified via NiNTA affinity chromatography. Ufd1-His and Npl4 were combined to allow heterodimer formation. After thawing, cell suspensions were supplemented with 20 mM imidazole, 1 mM PMSF and 1 mg/ml lysozyme. After stirring at 4 °C for 30 minutes, cells were lysed by sonication followed by centrifugation at 16.000g for 1 h. The soluble supernatant was filtered and loaded onto a HisTrap™ NiNTA affinity column (Cytiva). The column was washed with 200 ml lysis-buffer supplemented with 20 mM imidazole and eluted with 25 ml of lysis-buffer supplemented with 300 mM imidazole. Proteins were further purified by size exclusion chromatography using either a Superdex 75 (His-gp78-ubc7, His-Cys-Ub K48R) or Superdex 200 (Cdc48, Ufd1-His + Npl4, His-Ube1) gel filtration column (Cytiva) with gel filtration-buffer (50 mM HEPES pH 7.4, 150 mM KCl, 2 mM MgCl_2_, 5% glycerol), aliquoted and stored at -80 °C. Prior to gel filtration, His-SUMO-Cdc48 was incubated with His-SENP2 SUMO-protease, dialyzed overnight at 4 °C against gel filtration-buffer (+ 20 mM imidazole) and passaged through a HisTrap™ NiNTA column. Untagged wild-type ubiquitin was purified as previously described^57^.

#### Ubiquitination of substrates

Substrates (MBP-ubi and MBP-ubi-ubi-MBP, 10 µM) were incubated with His-Ube1 (2 µM), His-gp78-ubc7 (20 µM) and ubiquitin wt (100 µM) in ubiquitination buffer (50 mM HEPES pH 7.4, 150 mM KCl, 10 mM MgCl_2_, 5% glycerol, 10 mM ATP) at 37 °C. Ubiquitin was added stepwise over the first 3 hours before incubation overnight. The solution was passed through a MBPTrap column (Cytiva) to remove enzymes and unanchored ubiquitin chains. The column was washed with gel filtration buffer and eluted with gel filtration buffer supplemented with 20 mM maltose. Substrates were further purified by gel filtration using a Superose™ 6 increase SEC column (Cytiva). Fractions containing highly-ubiquitinated substrates were pooled, aliquoted, flash-frozen in liquid N_2_ and stored at -80°C.

Polyubiquitinated MBP-ubi was subjected to a second round of ubiquitination with His-cys-ubiquitin K48R. The sample was purified via affinity chromatography on an MBPTrap column (Cytiva) followed by a HisTrap column (Cytiva) to isolate substrates that carried both an MBP and a His-tag (**Extended Data Fig.1**).

The fluorescent unfolding substrate His-ubi-ubi-mEos3.2 was ubiquitinated as described above, except that the His-tags on Ube1 and gp78-ubc7 were removed before. This allowed separation of the ubiquitinated substrates from the enzymes post reaction through affinity purification with at HisTrap™ NiNTA affinity column (Cytiva). After subsequent gel filtration using a S200 16/600 column (Cytiva) fractions were separated depending on the length of the ubiquitin chains into long (L), medium (M) and short (S) (**Extended Data Fig. 1**).

### ATPase assay

The ATPase rate of Cdc48 was measured by combining Cdc48 (50 nM, hexamer) and Ufd1 + Npl4 (100 nM) with either no substrate, MBP-ubi-ubi-MBP (100 nM), or poly-ubiquitinated MBP-ubi-ubi-MPB (100 nM) in reaction buffer (50 mM HEPES pH 7.4, 100 mM KCl, 5 mM MgCl2, 1 mM DTT) and adding an ATPase regeneration system (1 mM ATP, 0.15 mM NADH, 3.75 mM phosphoenolpyruvate, 0.5 mM pyruvate kinase/lactic dehydrogenase mix [Sigma Aldrich]). Each condition was prepared in triplicate and the NADH absorbance at 340 nm was measured at 25 °C for 2 h. Linear absorbance decay was calculated (GraphPad Prism) and normalized to obtain the relative ATPase rates (**Extended Data Fig. 7**).

#### Labelling of Cdc48 and Ufd1+Npl4

The fluorescent dyes (NHS-Cy3B [Cytiva] for Cdc48; NHS-Atto643 [ATTO-TEC] for Ufd1+Npl4) were diluted in DMSO (10 mM) and added to 30 µM of the proteins in 1×PBS (pH set to 8.3 with 0.2 M sodium bicarbonate buffer pH 9.0) at 100 µM final concentration. After incubation over night at 4 °C, reactions were stopped by the addition of 1 mM Tris pH 8.0 and solutions were purified twice with Zeba spin desalting columns (Thermo Fisher Scientific) equilibrated with gel filtration buffer.

### Cdc48 binding experiment

The binding of Cdc48 to poly-ubiquitinated MBP-ubi-ubi-MBP was determined by MBP pulldown. Solutions containing different combinations of Cdc48-Cy3B (hexamer), Ufd1-His + Npl4 (Atto643), MBP-ubi-ubi-MPB and poly-ubiquitinated MBP-ubi-ubi-MBP (50 nM each) were incubated with amylose resin (New England Biolabs) for 1 hour at 4 °C in IP buffer (50 mM Tris pH 7.4, 150 mM KCl, 5 mM MgCl_2_, 5% glycerol, 1% Triton X-100, 2 mM β-mercaptoethanol). Beads were washed and eluted with 1.5× SDS-sample buffer at 95 °C for 5 minutes. Results were analysed by fluorescence scans with a Chemostar Fluorescence Imager (Ex.: 535 nm, Em.: 607 nm, Intas) and Odyssey Imager (Ex.: 685 nm, Em.: 730 nm, Licor) and finally by Coomassie staining (**Extended Data Fig. 8**).

### Mass Photometry

The length of ubiquitin chains attached to the substrates was determined using a TwoMP mass photometer (Refeyn). Samples were measured at 24 nM dilution following manufacturer instructions. The number of attached ubiquitins was calculated by subtracting the mass of the base substrate without ubiquitin (50 kDa (ubi-MBP) or 100 kDa (MBP-ubi-ubi-MBP) respectively) and dividing by 8.5 kDa (**Extended Data Fig. 2)**.

### Deubiquitination assay

Poly-ubiquitinated substrate (5 µg) was incubated with 1 µM of deubiquitinating enzyme (YOD1 or Ataxin-3) for 30 minutes at 37°C, followed by SDS-PAGE analysis (**Extended Data Fig. 3).**

### Fluorescent unfolding assay

Fluorescent unfolding substrate (His-ubi-ubi-mEos3.2) was irradiated with a longwave UV-lamp (UVP Blak-RayTM B-100AP) for 2 h, to induce photoconversion green (Ex.: 500 nm, Em.: 520 nm) to red (Ex.: 540 nm, Em.: 580 nm) fluorescence. Photoconverted Eos (red) is unable to refold after unfolding by Cdc48, allowing it to be used as reporter for unfolding activity. Substrates with different degrees of ubiquitination (L, M, S) were mixed at 125 nM concentration together with Cdc48 (175 nM hexamer) and Ufd1-Npl4 (500 nM) in unfolding buffer (25 mM HEPES, pH 7.4, 100 mM KCl, 5 mM MgCl2, 5 % glycerol, 1 mM DTT) in incubated at 30°C for 5 minutes. The unfolding reaction was started by addition of ATP to 2 mM and red mEos3.2 fluorescence was monitored for 40 minutes using a Cary-Eclipse fluorescence-spectrophotometer (Varian, E.: 540 nm, Em.: 580 nm). Results were plotted with GraphPad Prism (**Extended Data Fig. 1C-D)**.

### Protein-DNA construct preparation

In order to eliminate reducing agents, the ubiquitinated protein constructs underwent buffer exchange using a PD-10 desalting column (GE Healthcare). 20-base pair (bp) maleimide single-stranded DNA oligos (Biomers) were linked to terminal cysteines of the two constructs (polyubiquitinated MBP-ubi-ubi-MBP and MBP-ubi) for 1 hour at room temperature in TCPOC coupling buffer (100 mM sodium phosphate, 150 mM NaCl, 10 mM EDTA, pH 7.2). Uncoupled oligos were removed using amylose resin (New England Biolabs) as described above for substrate purification. 1.3 kbp DNA handles containing a 20-nucleotide single-stranded overhang complementary to the protein-linked DNA oligos were produced via polymerase chain reaction using the pUC19 plasmid (New England Biolabs) as the template, as previously reported *(27).* The side of the handles not containing the overhang was labelled with either double digoxigenin or biotin (approximately 50% each). The protein-oligo complex was then ligated to the DNA handles using T4 ligase (New England Biolabs) for 6 h at 16°C followed by overnight incubation on ice.

### Single-molecule assays

#### Optical tweezers setup

The single-molecule experiments were performed using a C-Trap (Lumicks). In brief, the instrument consists of two optical traps formed by a single high-intensity, polarisation-stable 1064 nm laser split into two orthogonally-polarised beams. Samples are manipulated in a monolithic laminar flow cell with five separate flow channels controlled by a passive pressure-driven microfluidic system.

#### Sample preparation

Protein-bead samples were prepared daily by combining the protein-DNA construct (80 μM, 1 μL) with Ø 2 μm anti-digoxigenin-coated polystyrene beads (0.1% w/v, 1 μL; Spherotech) in a sample buffer (25 mM HEPES pH 7.4, 100 mM KCl, 5 mM MgCl_2_; 10 μL). Samples were incubated overhead for 25 min at 4°C, then made up to a final volume of 350 μL with sample buffer, prior to measurement. NTV-bead samples were prepared by diluting Ø 2 μm Neutravidin-coated polystyrene beads (0.5% w/v, 0.2 μL; Spherotech) in sample buffer (350 μL).

A measurement buffer was prepared by supplementing the sample buffer with an oxygen scavenging system (50 mM glucose, 3 units mL^-1^ pyranose oxidase [Sigma-Aldrich], 90 units mL^-1^ catalase [Sigma-Aldrich])^58^, ATP regeneration system (1 mM ATP / ATPγS, 3 mM phosphoenol pyruvate [Sigma-Aldrich], 20 mg mL^-1^ pyruvate kinase [Sigma-Aldrich]), TCEP-HCl (1.25 mM), Cdc48 (500 μM, in the +Cdc48 condition only), and Ufd1-Npl4 (1 mM, in the +Cdc48 condition only).

#### Optical tweezers operation

During experiment, a protein-bead was collected from one laminar flow channel in one trap, and an NTV-bead from a second channel in the other. The beads were moved to a third channel containing the measurement buffer prior to tether formation. The beads were repeatedly brought into close proximity and back until a slight increase in measured force upon retraction indicated formation of a tether. Single tethers were identified according to three criteria: the tether length (*ca.* 0.9 µm corresponding to the two 1.3 kbp dsDNA handles); presence of the characteristic DNA overstretching motif at 65 pN^59,60^; and presence of the characteristic unfolding transition(s) of MBP^61,62^ (1-2 molecules, depending on the construct – **Extended Data Fig. 6**).

For the ubiquitin unfolding experiments (**Figure 1J-L**), measurements were taken in a cycling ‘force spectroscopy mode: the steerable trap was moved at a constant rate of 0.1 µm s^-1^ between a minimum trap separation of 0.8 µm and a maximum applied force of 65 pN repeatedly until tether breakage.

All other measurements were taken in ‘constant-distance’ mode: the steerable trap was moved at a constant rate of 0.1 µm s^-1^ until the applied force reached 10 pN. The traps were then held stationary for a period of 500 - 1000s, or until tether break.

