## Extended Data for "Direct observation of ATP-driven ubiquitin chain handling by Cdc48"

Extended Data Figures 1 to 15

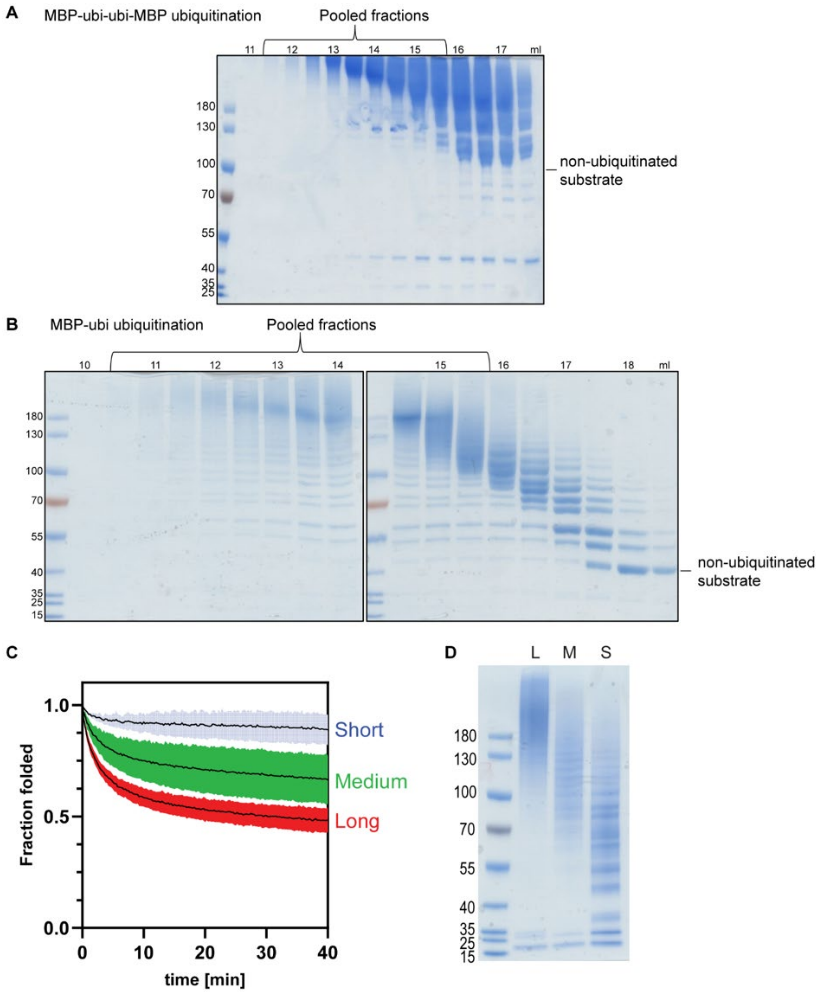


**Extended Data Fig. 1. Ubiquitination. A,** Ubiquitination for the conformationally free construct (Figure 2A-B)**.** Substrates were ubiquitinated with untagged ubiquitin, using the E1 enzyme Ube1 and a chimeric fusion of the E2 enzyme ubc7 and the E3 enzyme gp78 which produces K48-linked chains. Reaction products were separated by size exclusion chromatography (Superose 6, Cytiva) and highly ubiquitinated fractions were pooled. **B,** Ubiquitination for the tethered ubiquitin chain construct (Figure 1A-B). See panel A text and Methods for further details. **C,** Fluorescent substrates His-ubi-ubi-Eos (poly-ub) were prepared with three different degrees of ubiquitination (long L, medium M, short S) and unfolded by Cdc48-Ufd1-Npl4 after addition of ATP. Comparison of the loss of Eos fluorescence over time shows that long ubiquitin chains are most efficient in unfolding (n = 3, mean and SEM). **D,** SDS-PAGE coomassie stain of His-ubi-ubi-Eos (poly-ub) with long (L), medium (M) and short (S) ubiquitin chains. The substrate was ubiquitinated with the gp78-ubc7 fusion enzyme, attaching K48 linked ubiquitin chains, which were then sorted into long, medium and short chain length by gel filtration.

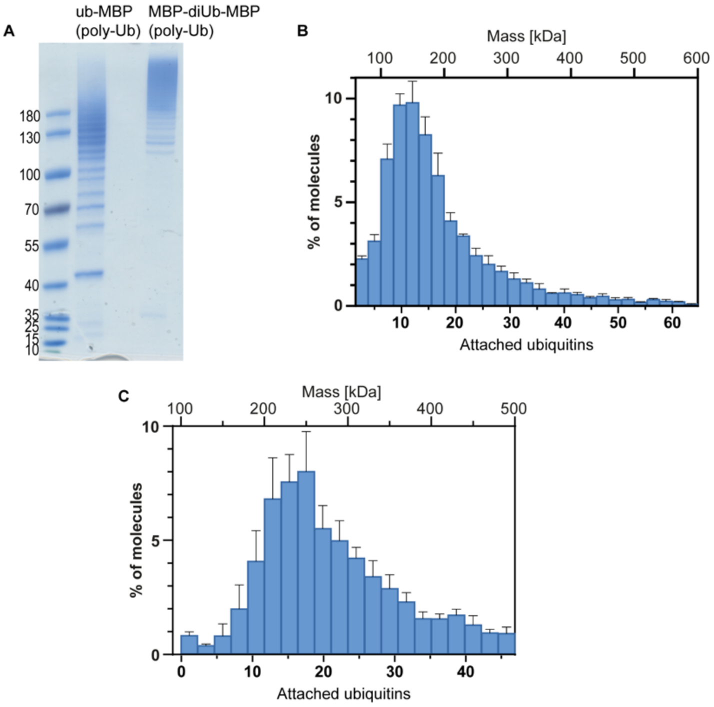


**Extended Data Fig. 2. Measurement of ubiquitin chain lengths.** **A**, SDS-PAGE coomassie stain of ub-MBP and MBP-ubi-ubi-MBP (ubiquitins are poly-ubiquitinated). **B**, Mass photometry measurement of ubiquitin-chain construct ub-MBP (poly-ub). Results from three independent measurements showing the mass distribution as % of all molecules in the sample (means and SEMs). The number of attached ubiquitins was calculated based on the mass of the base substrate (50 kDa) and the mass of one ubiquitin (8.5 kDa). **C**, Mass photometry measurement of MBP-ubi-ubi-MBP (poly-ub). Results from three independent measurements showing the mass distribution as % of all molecules in the sample (means and SEMs). The number of attached ubiquitins was calculated based on the mass of the base substrate (100 kDa) and the mass of one ubiquitin (8.5 kDa).

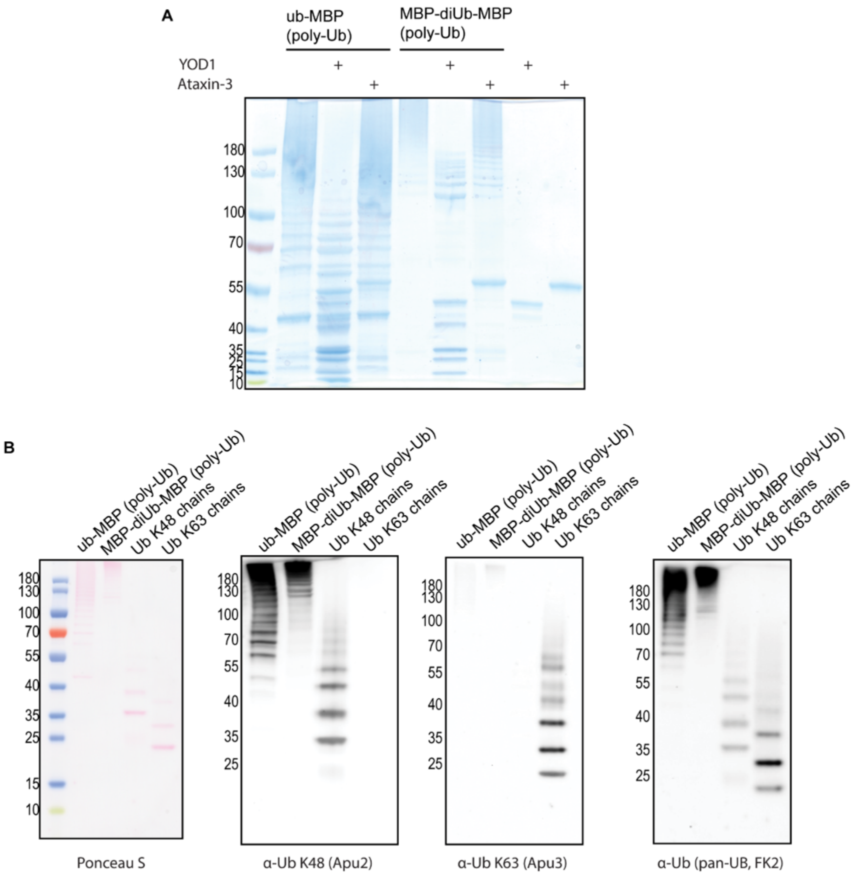


**Extended Data Fig. 3. Ubiquitin linkage type of the substrates**. **A**, Deubiquitination assay of ub-MBP and MBP-diUb-MBP (ubiquitins are poly-ubiquitinated) with the deubiquitinating enzymes YOD1 (K48 specific) and Ataxin-3 (K63 specific). Only YOD1 is capable of digesting the attached ubiquitin chains. **B**, Western-blot analysis of ub-MBP (poly-ub) and MBP-ubi-ubi-MBP (poly-ub) with chain type specific ubiquitin antibodies, compared to purchased chains (3-7 ubiquitin molecules) with distinct ubiquitin linkages. All samples can be detected by Ponceau S stain and a pan-ubiquitin antibody (FK2), but substrates can only be detected with K48- and not with K63-specific antibodies.

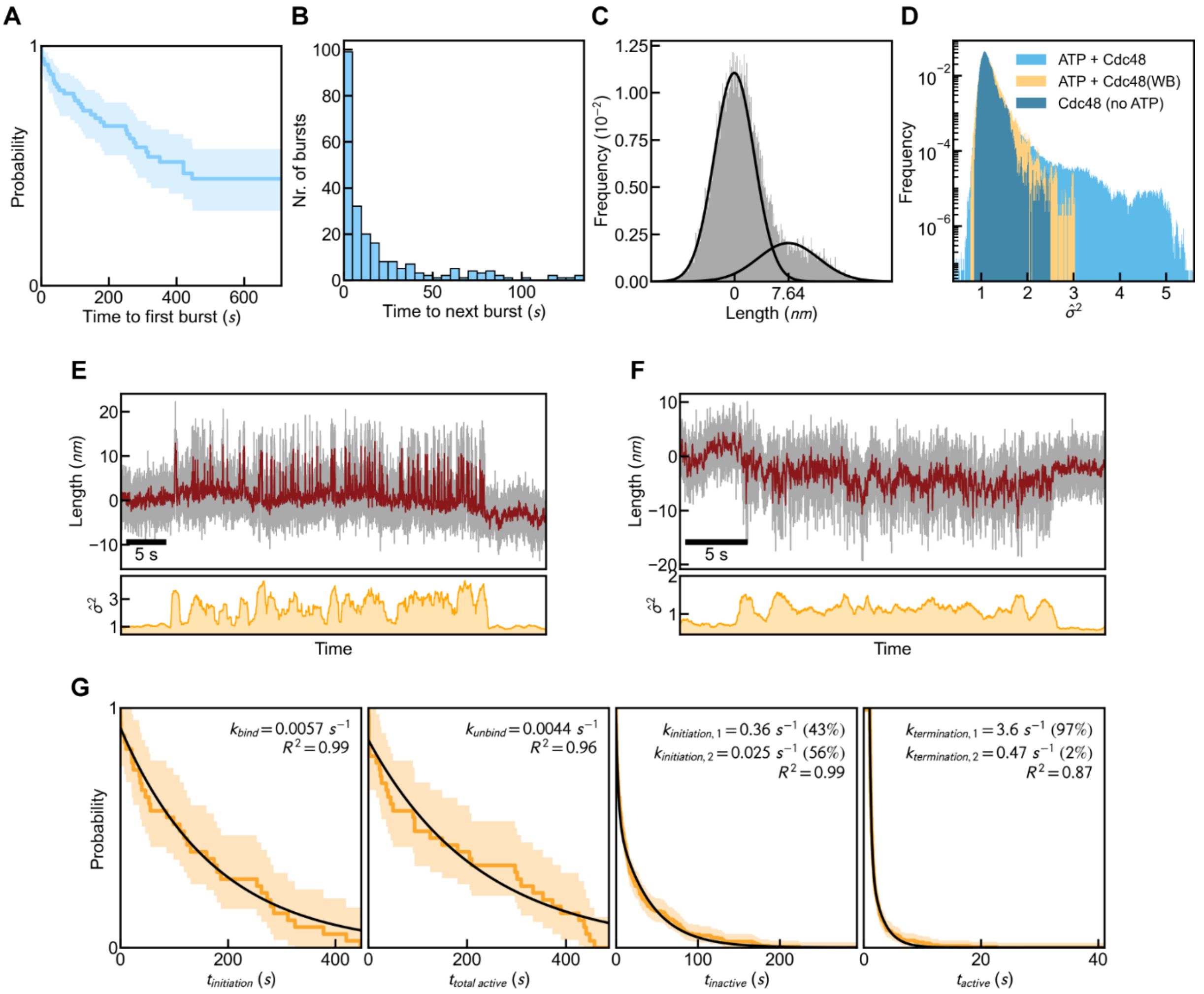


**Extended Data Fig. 4. Non-Processive activity – burst intervals, waiting time, and duration. A,** Histogram of the time interval in between consecutive bursts of activity (**Figure 1D-E**), for the tethered ubiquitin chain construct (**Figure 1A-C**). The distribution shows that this time interval is short, of order seconds, indicating that one burst of activity is followed quickly afterwards by another. N=5 events with t > 150 s are truncated for clarity. **B,** Survival analysis for the waiting time until a non-processive activity (**Figure 1D-E**) is observed after starting the experiment, for the tethered ubiquitin chain construct (**Figure 1A-C**). The Kaplan-Meier curve indicates the fraction of experiments that do not (yet) show non-processive activity, as a function of elapsed time in the experiment. **C,** Histogram of length measured during the non-processive burst shown in **E.** A Gaussian Mixture fit to the histogram gives two peaks separated by 7.64 nm, the average fluctuation length during a burst. **D**, Distribution of $\hat{\sigma}^2$ that quantifies the fluctuation intensity in the presence of Cdc48 with ATP (blue), Walker B mutant E315Q with ATP (yellow) and Cdc48 with no ATP (dark blue). **E, F,** Top: length vs. time in presence of Cdc48 and ATP, showing non-processive Cdc48 activity, for the tethered ubiquitin chain construct (**Figure 1A-C**). Raw and filtered signals are shown in grey and red respectively.

Bottom: $\hat{\boldsymbol{\sigma}}^2$-time trace, showing the non-processive bursts as periods of increased fluctuation intensity above the base level. The length fluctuations can start either from the higher (D) or the lower (E) level. The data show that bursts can persist for tens of seconds. **G.** Survival functions of four time coordinates extracted from the automated detection of non-processive action by Cdc48. From left to right, the time coordinates are: the time between the beginning of the measurement and the first detected event, $t_{initiation}$; the time between the first and last detected events, $t_{total\ active}$; the time between detected events, $t_{inactive}$, and the duration of detected events, $t_{active}$. Each survival function is defined using a Kaplan-Meier estimator, and the decay rate(s) determined by an exponential fit to each function. The rate(s) k and coefficient of determination $R^2$ for each fit is annotated top-right. Exponential fits to each survival function respectively yield rate constants $k_{bind}$ (rate of observing first activity, which may include binding of Cdc48 to substrate), $k_{unbind}$ (rate of switching to full activity arrest, which may include Cdc48 unbinding), $k_{initiation}$ (rate of switching from inactive to active), and $k_{termination}$ (rate of switching from active to inactive). The measured $k_{bind}$ and $k_{unbind}$ match those obtained for the processive activity of Cdc48 (**Extended Data Fig. 12E**). The measured $k_{initiation}$ is much greater than $k_{bind}$, consistent with the multiple bursts of activity identified after initial binding of Cdc48 to the substrate.

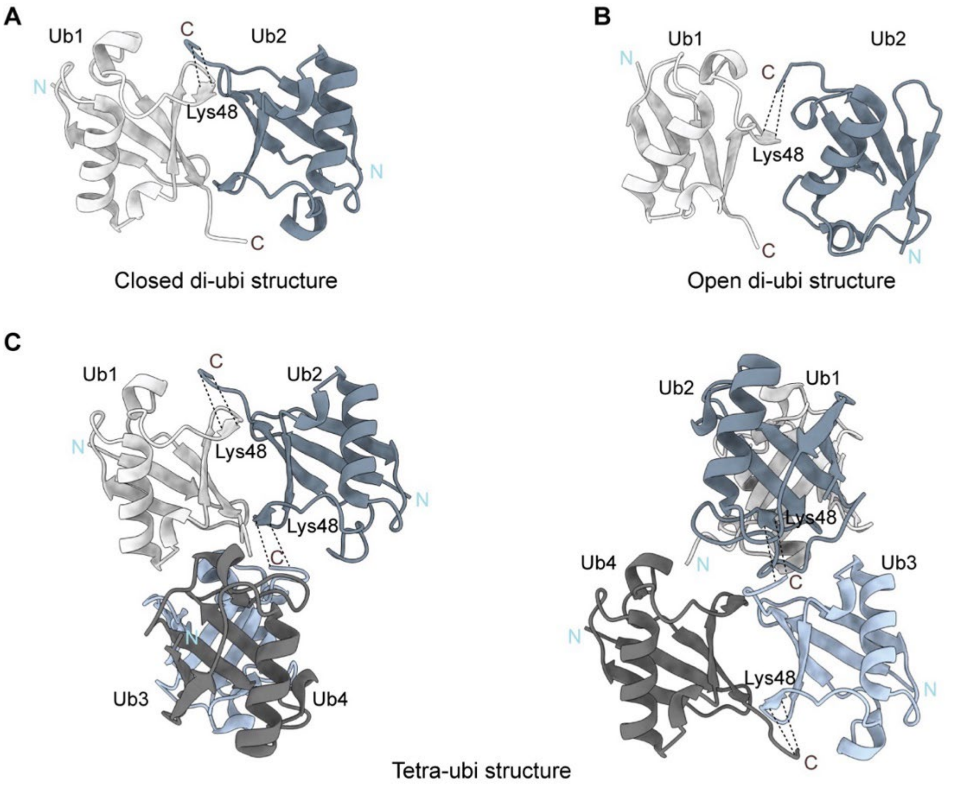


**Extended Data Fig. 5. Structural diversity of Lys-48 linked ubiquitin dimers and tetramers. A,** Closed structure of lys-48 linked di-ubiquitin (PDB 3M3J)[54]. **B,** Open structure of lys-48 linked di-ubiquitin structure (PDB 3AUL)[55]. These differences in conformation (panels A and B) were shown to affect di-ubiquitinase activity. Transitioning between the closed (A) and the open (B) conformations causes a length change of 0.77 nm between the Gly-75 residue (the C-terminus) of Ub1 and Lys-48 residue of Ub2. In Figure 1H (middle) we speculate about remodeling of polyubiquitin, involving the 3 folded ubiquitin moieties bound to Ufd1-Npl4. Open-closed transition for two ubiquitin-ubiquitin interfaces would give a measured length change of about 1.5 nm. Note that here we consider remodelling induced by Cdc48, which is restricted to 3 ubiquitin moieties. **C,** Compact structure of lys-48 linked tetra ubiquitin structure (PDB 2O6V)[22], showing the interactions between ubiquitin moieties, for instance between $Ubi_1$ and $Ubi_4$. The tetra-ubiquitin structure resembles a pair of stacked di-ubiquitin, and may also exhibit structural transitions. How the alternative conformations of polyubiquitin are populated in time could well depend sensitively on bound factors, as well as in the tethered ubiquitin construct (**Figure 1**), which in turn is counter-selected by the non-processive Cdc48 activity, thus blocking transition to processive Cdc48 translocation. In contrast, the conformationally free construct is shown to transition to processive translocation (**Figure 2**).

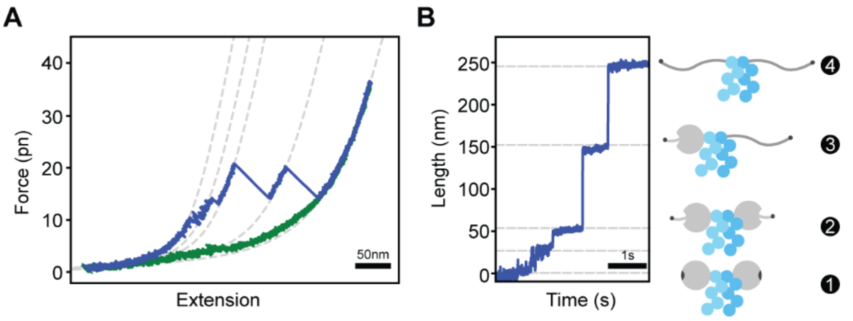


**Extended Data Fig. 6. Mechanical manipulation of MBP-Ubiquitin-Ubiquitin-MBP.** Force-extension (**A**) and contour length-time (**B**) plots showing substrate unfolding. The force extension curve displays a saw-tooth pattern, reflecting the unfolding of two MBP molecules within the MBP-ubi-ubi-MBP construct. Unfolding begins with the initial unravelling of α-helical structures (20-25 nm each, 1 to 2), followed by two sudden length increases (2 to 3 and 3 to 4) of approximately 100 nm each, corresponding to unfolding of the MBP core. Protein contour length increases corresponding to these unfolding events, as determined using the worm-like chain (WLC) model (Methods). "Length" indicates the contour length of the unfolded part of the protein, with the folded part having negligible effect on the measurement. Dotted grey lines represent the WLC-fits applied to the data. Note that ubiquitin is a very stable molecule and typically does not unfold.

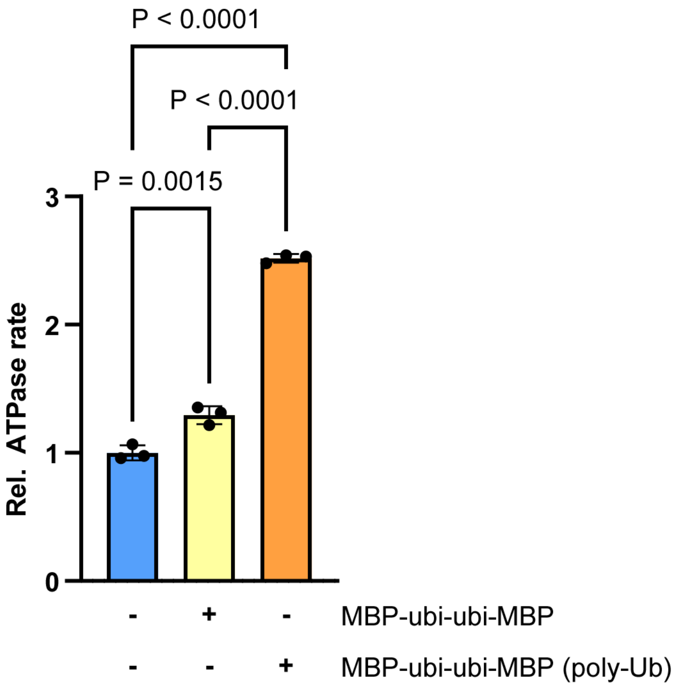


**Extended Data Fig. 7. Poly-ubiquitinated MBP-ubi-ubi-MBP stimulates the ATPase rate of Cdc48.** The ATPase rate of Cdc48 in the presence of Ufd1-Npl4 was calculated based on the NADH oxidation rate in an ATP regenerating system (ATP, NADH, phosphoenolpyruvate, pyruvate kinase, lactate hydrogenase) by measuring the absorbance at 340 nm. Addition of MBP-ubi-ubi-MBP only results in small increase of the ATPase rate (~20 %), while the addition of poly-ubiquitinated MBP-ubi-ubi-MBP results in a significant increase by a factor of ~2.5, showing that the latter is being processed by Cdc48.

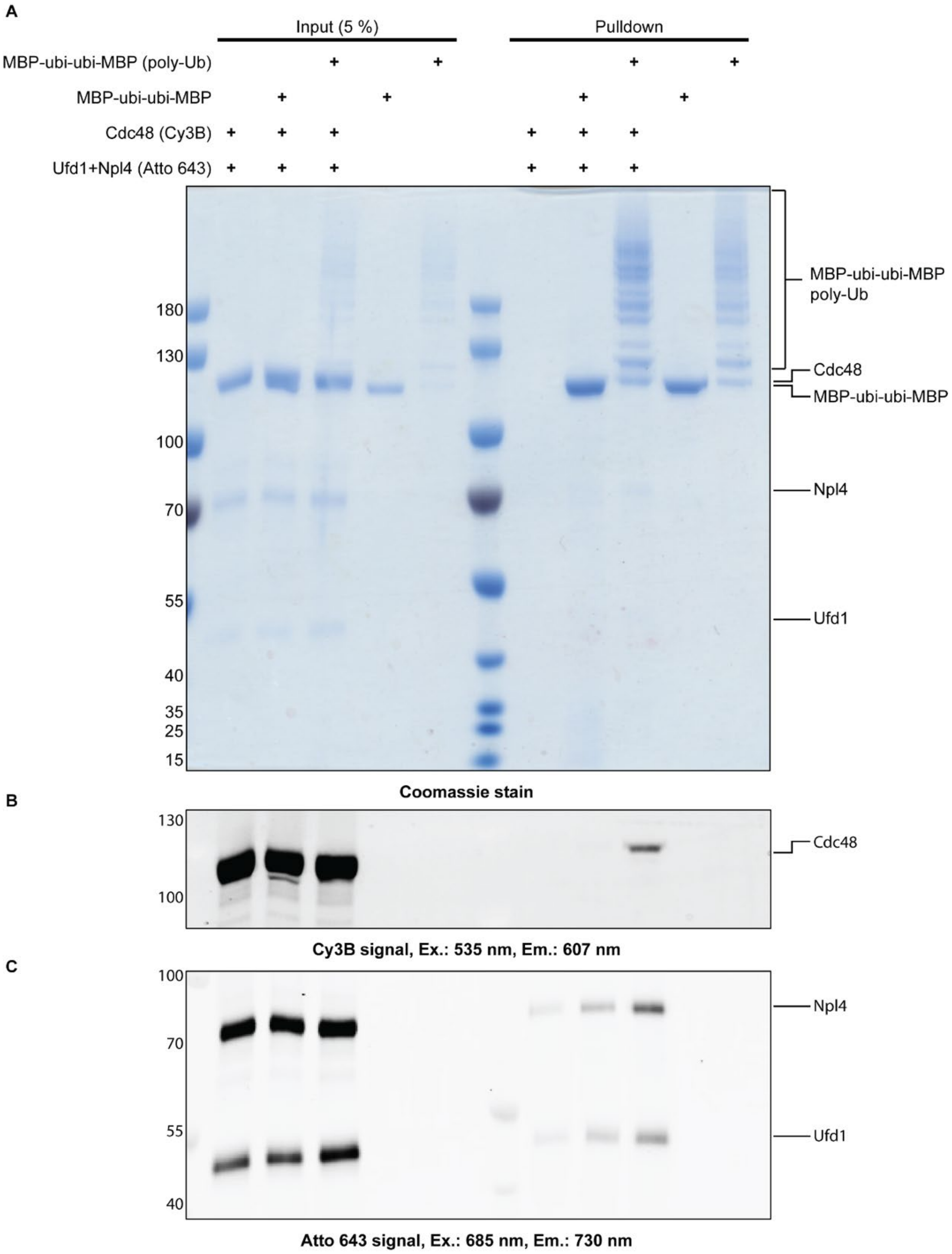


**Extended Data Fig. 8. Interaction between MBP-ubi-ubi-MBP and Cdc48.** Binding of MBP-ubi-ubi-MBP to Cdc48 (labelled with Cy3B) and Ufd1-Npl4 (labelled with (Atto 643) requires poly-ubiquitination. Pulldown of MBP-ubi-ubi-MBP or polyubiquitinated MBP-ubi-ubi-MBP with amylose resin shows that only the poly-ubiquitinated substrate interacts with Cdc48 and Ufd1-Npl4, based on Coomassie staining (**A**), Cy3B signal (Ex.: 535 nm, Em.: 607 nm) (**B**) and Atto 643 signal (Ex.: 685 nm, Em.: 730 nm) (**C**).

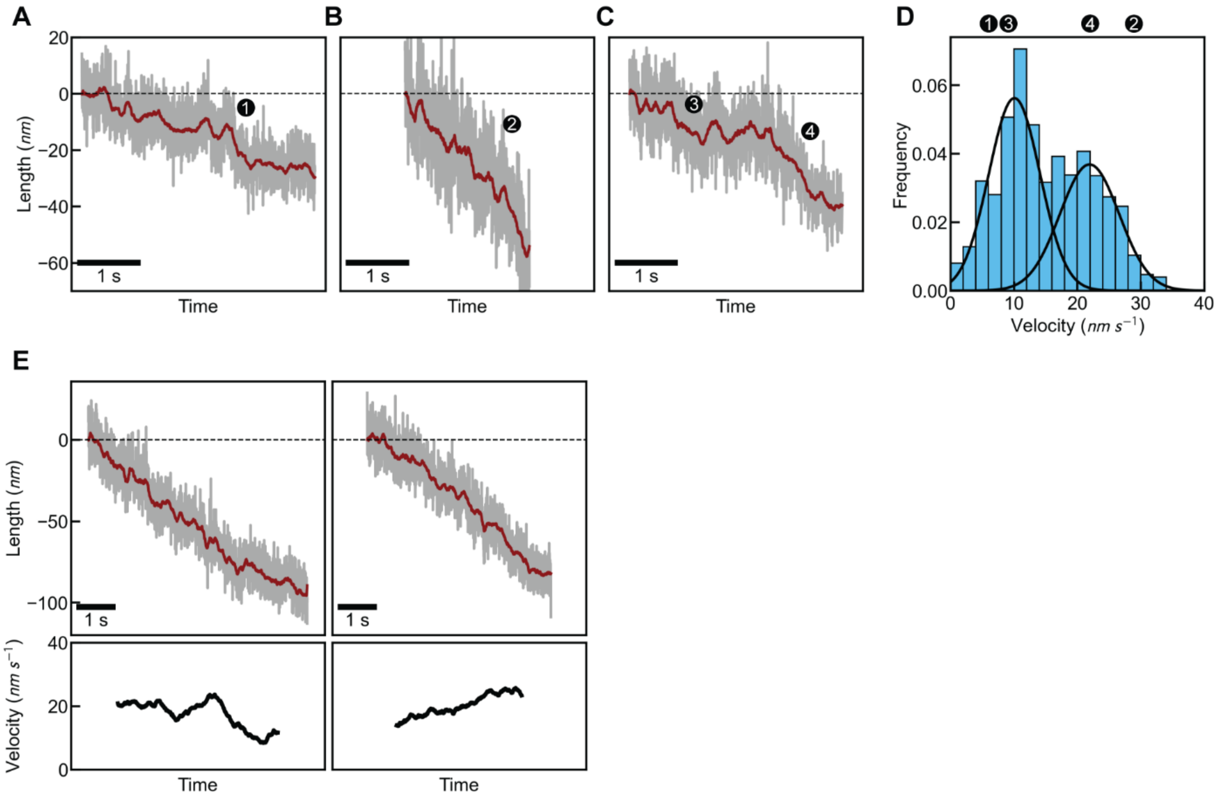


**Extended Data Fig. 9. Translocation velocity – diversity and continuity in time. A-C,** length vs. time in presence of Cdc48 and ATP, showing the diverse translocation velocities during processive Cdc48 action, for the conformationally-free construct (**Figure 2A-B**). Examples indicate comparatively slow (A), fast (B), and switching between slow and fast translocation in time (C). The data are consistent with Cdc48 translocating one arm of an inserted polypeptide loop (slow) and two arms simultaneously (fast). **D,** Velocity distribution of all translocation events, with numbers corresponding to the examples shown in panels A-C. **E,** Translocation speed stability. Top: length vs. time in presence of Cdc48 and ATP. Bottom: corresponding translocation velocity versus time in a 1.5-second moving window. The data show that translocation velocity remains roughly constant in time, which is a hallmark of processive activity.

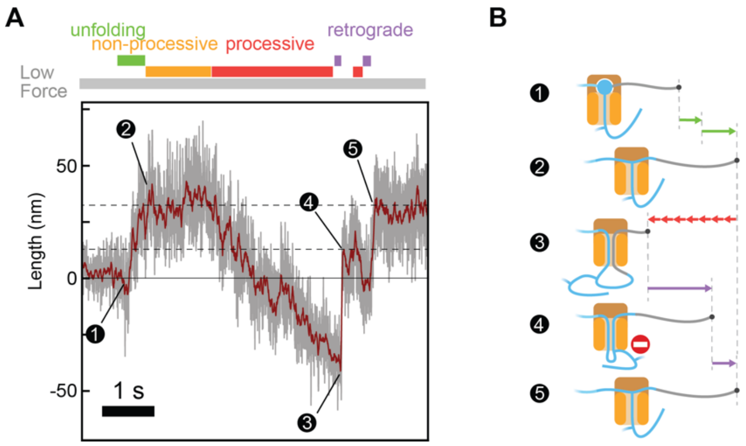


**Extended Data Fig. 10. Complete retrograde movement. A,** Length vs. time in presence of Cdc48 and ATP for the conformationally-free construct (**Figure 2A-B**). The data show a number of subsequent events: Ubi$_0$ unfolds (1); a period of non-processive activity (2); processive translocation (ending at 3), retrograde movement in two steps (3 to 5). The length at which the transient retrograde movement arrest (4) occurs is consistent with the branch point (in *trans*) acting as a barrier against immediate *cis*-release. **B,** Cartoons illustrating these five phases.

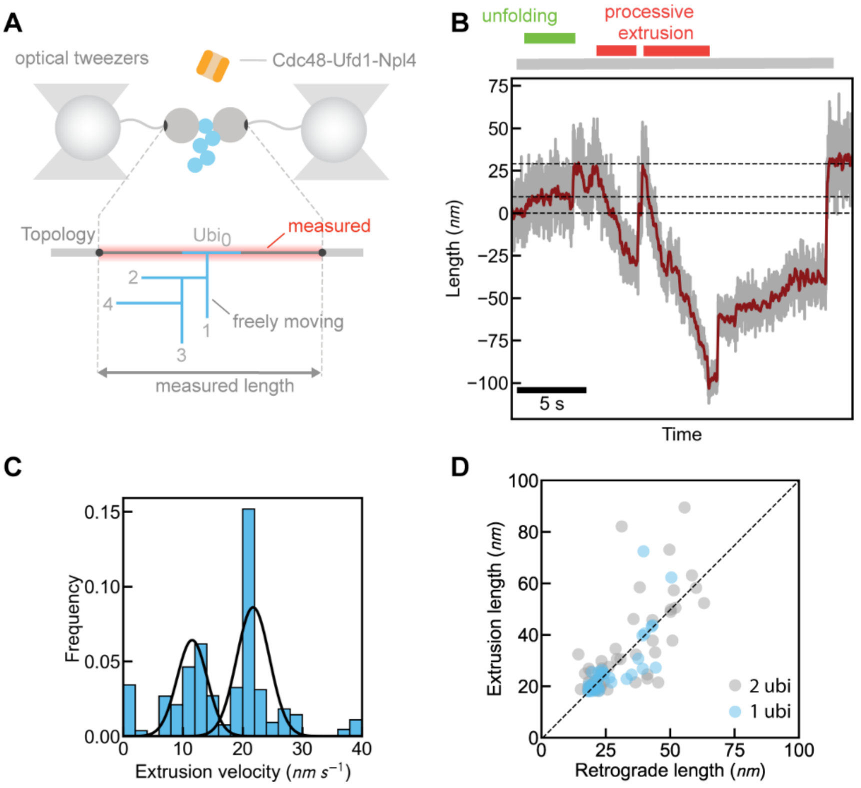


**Extended Data Fig. 11. Processive extrusion on single polyubiquitin chain construct. A**, Optical tweezers assay for conformationally-free construct with only one ubiquitin chain. **B**, Extrusion event (Construct length vs. time) with Cdc48, showing processive Cdc48 activity. **C**, Distribution of extrusion velocity. **D**, Extruded length vs. retrograde lengths for the 1 ubi (blue) and 2 ubi (grey) construct.

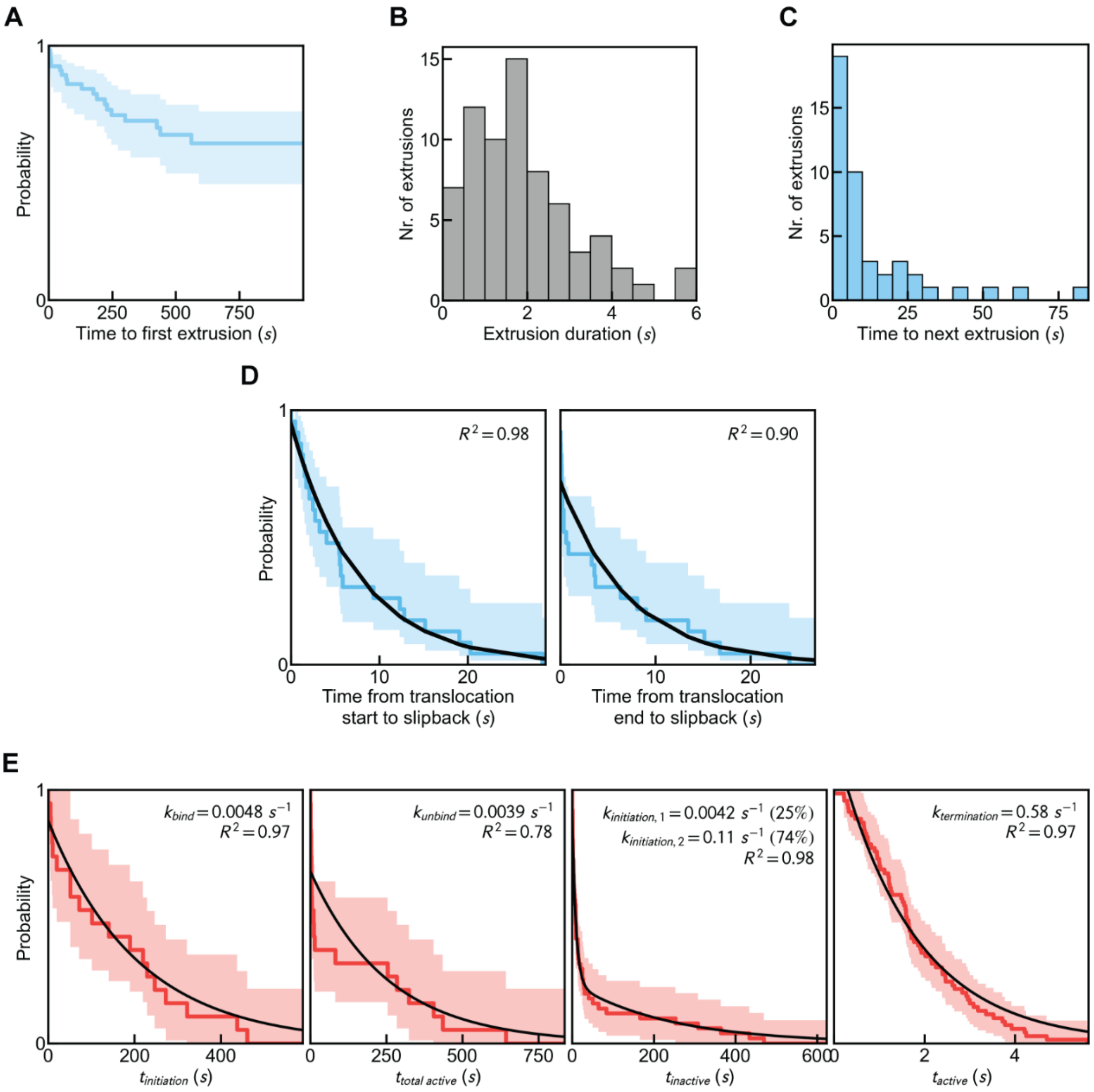


**Extended Data Fig. 12. Translocation initiation, duration and repeated activity. A,** Survival analysis for the waiting time until a processive extrusion activity (**Figure 2C**) is observed after starting the experiment, for the ubiquitin-free construct (**Figure 2A-B**). The Kaplan-Meier curve indicates the fraction of experiments that do not (yet) show processive activity, as a function of elapsed time in the experiment. The plateau level shows that processive translocation events are detected in 26.7% of measured molecules, while the region where the curve decreases indicates that the waiting time is broadly distributed up to hundreds of seconds. Note that specifically the processive translocation of Ubi$_0$ and MBP is detected in this construct. Hence, the initial binding Cdc48 to the initiator ubiquitin (Ubi$_{init}$) and the subsequent processing of the ubiquitin moieties Ubi$_{init-1}$, Ubi$_{init-2}$, … down to Ubi$_1$ is not observed directly and rather occurs during this waiting time. The length of the ubiquitin chain can be more than 8 ubiquitin monomers. **B,** Histogram of the duration of observed processive translocation events, showing processive activity lasts for seconds. **C,** Histogram of the elapsed time between consecutive translocation events. These pauses in activity generally follow a retrograde movement that ends the first translocation event. These data show that restarting translocation takes several seconds. N=7 events with t > 150 s are truncated for clarity. **D,** Survival analysis of time between translocation start and retrograde movement (left), and time between the end of translocation and retrograde movement (right). **E**, Survival functions of four time coordinates extracted from the automated detection of processive translocation by Cdc48. From left to right, the time coordinates are: the time between the beginning of the measurement and the first detected event, $t_{initiation}$; the time between the first and last detected events, $t_{total\ active}$; the duration of detected events, $t_{active}$; and the time between detected events, $t_{inactive}$. Each survival function is defined using a Kaplan-Meier estimator, and the decay rate(s) determined by an exponential fit to each function. The rate(s) k and coefficient of determination $R^2$ for each fit is annotated top-right. Exponential fits to each survival function respectively yield rate constants $k_{bind}$ (rate of observing first activity, which may include binding of Cdc48 to substrate), $k_{unbind}$ (rate of switch to full activity arrest, which may include Cdc48 unbinding), $k_{initiation}$ (rate of switching from inactive to active), and $k_{termination}$ (rate of switching from active to inactive). The measured $k_{bind}$ and $k_{unbind}$ closely match those obtained for the non-processive action of Cdc48 (**Extended Data Fig. 4G**). The $k_{initiation}$ is much greater than $k_{bind}$, consistent with the multiple rounds of translocation and slipback identified after initial binding of Cdc48 to the substrate.

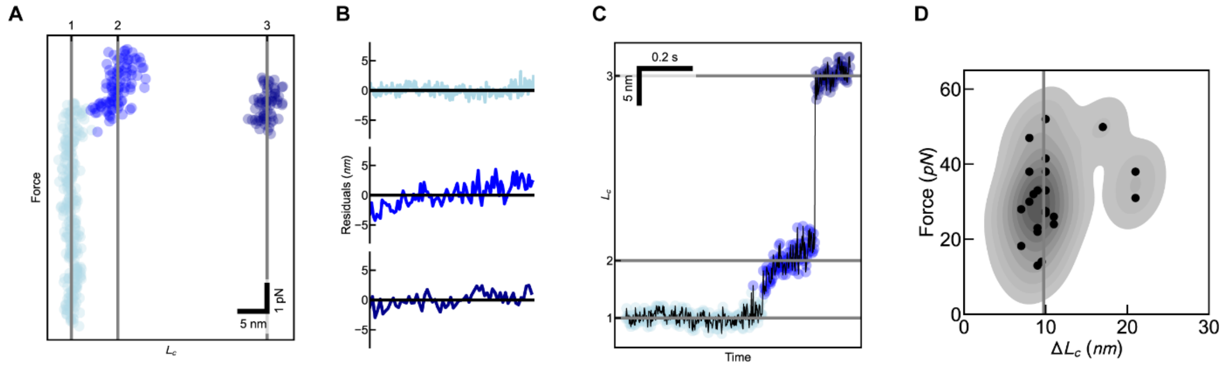


**Extended Data Fig. 13. Unfolding analysis. A)** After worm-like chain fitting of measured Force-Extension data, folding states are identifiable in the Force-$L_c$ domain as clusters with constant $L_c$. These states are identified algorithmically by K-means clustering. **B)** The quality of the worm-like chain fit to each state is assesed using a two-tailed Kolmogorov-Smirnov test of the fit residuals against a Gaussian distribution of zero mean[5]. Here, there is no statistical evidence to reject the fits at a 1% confidence level. **C)** After clustering, the folding states can be mapped back into the $L_c$-time domain. Unfolding lengths are measured as the differences between the median $L_c$ values between consecutive state (here 7.25 nm and 23.31 nm). **D)** Measured unfolding forces and lengths for the first unfolding step of ubiquitin, showing an average value of 10.4 +/- 3.8 nm.

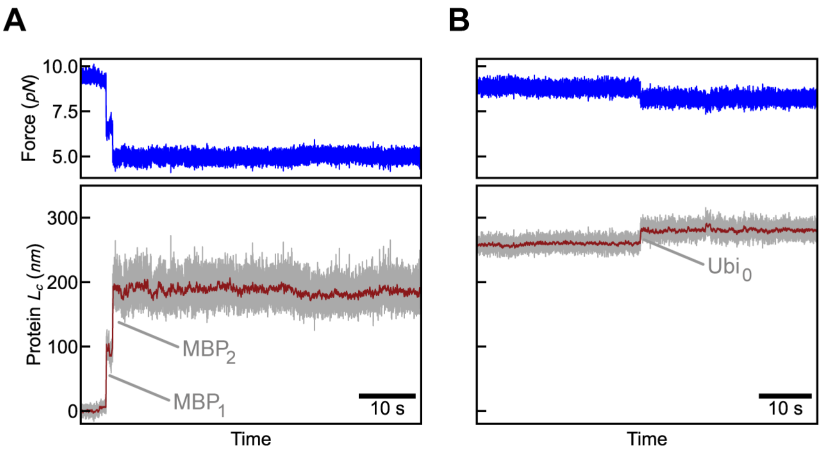


**Extended Data Fig. 14. MBP unfolding at low force. A)** Example data showing length for 2 MBP unfolding in two steps (bottom) when initially held at about 10 pN. As the trap positions are constant, the force correspondingly decreases to about 5 pN (top). In our typical Cdc48 activity measurements the applied force would be restored to about 10 pN after MBP unfolding to limit measurement noise. **B)** Example data showing measured length for one ubiquitin moiety unfolding (bottom) when initially held at about 9 pN. Corresponding Force measurements showing decreases by about 1 pN, owing to the constant trap distance and somewhat increased length (top).

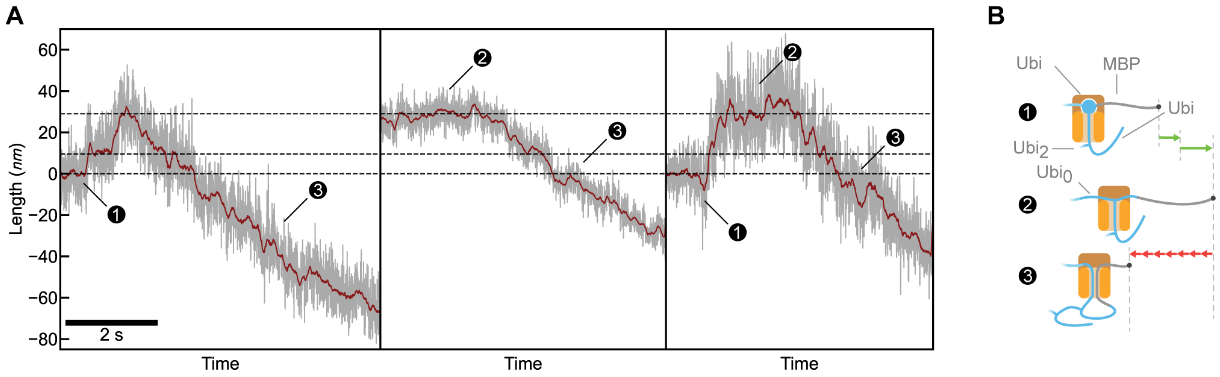


**Extended Data Fig. 15. Initial phases of translocation. A)** For comparison, here we display three example traces of Length vs Time at the start of Ubi0 translocation side by side. **B)** corresponding cartoons. When Ubi0 is unfolded (2), the first length decreases indicate branch point translocation through the Cdc48 pore. Sometime later, the branch point emerges in *trans* and continued translocation of the polypeptide loop (3) leads to translocation of the unfolded MBP substrate. The data does not show clear and significant differences in speed for branch point translocation compared to subsequent polypeptide loop translocation.
