## Supplementary Information for "Direct observation of ATP-driven ubiquitin chain handling by Cdc48"

***Data processing & statistical analysis***

For the ubiquitin unfolding experiments, folding events were identified and quantified using a semi-automated K-means clustering algorithm. All other results are based on custom detection algorithms for processive extrusion, retrograde movement, and non-processive action events, described in detail below. All calibration, fitting, and event identification was performed using custom scripts in Python. Statistical significance values, where reported, were determined using the Mann-Whitney U test (****: $p \leq 0.0001$, ***: $p \leq 0.001$, **$p \leq 0.01$, *$p \leq 0.05$).

*Data collection and pre-processing*

Force-distance data were collected at 50 kHz and decimated to 500 Hz prior to analysis. For each bead pair, the optical traps were calibrated by fitting a Lorentzian function to the power spectrum of the Brownian motion of the trapped beads[63]. The trapping laser intensity was kept constant for all measurements. To obtain the protein contour length, each force-extension curve was fit with two worm-like chain (WLC) models in series: the Odijk approximations for an extensible and inextensible WLC[64] for the DNA and protein components respectively.

For all time traces shown in main figures, data in grey are the raw data after decimation to 500 Hz. Data in red are filtered using a Savitzky-Golay filter[65] of 1st order, with a window of 51 points. Filtered data in figures is provided for illustrative purposes only.

*Quantification of signal variance in time*

Quantification of non-processive activity is based on measurement of the variance in the length signal in time. To obtain a good estimate, the variance of a time series is measured in a moving window across the length-time trace. For a length-time trace defined by vectors $\boldsymbol{L}$ and $\boldsymbol{t}$, the variance $\sigma_{n,i}^2$ at each time $t_i$ is given by:

$$\sigma_{n,i}^2 = \frac{1}{n}\sum_{j=i-n/2}^{i+n/2} \left(L_j - \mu_i\right)^2$$

where $n$ is the size of the moving window and $\mu_i$ is the mean value within the window, given by:

$$\mu_i = \frac{1}{n}\sum_{j=i-n/2}^{i+n/2} L_j$$

Signal variance is dependent on a number of environmental factors, including local temperature, bead size and trap strength. To facilitate comparison between measurements, a 'baseline' variance ($\sigma^2_{baseline}$) is taken as the median of the lowest 20 % of computed variance values. The variance vector is then normalised by this baseline as:

$$\hat{\boldsymbol{\sigma}}^2_n = \frac{\boldsymbol{\sigma}^2_n}{\sigma^2_{baseline}}$$

After normalisation, data without Cdc48 action have $\hat{\boldsymbol{\sigma}}^2_n = 1$, while values significantly above 1 indicate non-processive activity by Cdc48. In all main figures, the normalised variance over a 1 s moving window is used. For simplicity, the $n$ subscript is removed in main figures, leaving the normalised variance denoted $\hat{\boldsymbol{\sigma}}^2$.

*Non-processive activity detection*

Non-processive activity appears as a sharp increase in the measured variance of the length signal which does not correlate with unfolding or refolding of the tethered protein.

In the contour length domain (labelled "Length" in all main figures), unfolding and refolding transitions appear as discrete steps (**Extended Data Fig. 6**). Folding events may therefore be identified using AutoStepfinder, an algorithm optimised specifically for automated identification of step transitions in single-molecule time-series data[66]. At each folding state (between a pair of folding events), the baseline variance is re-computed using the method described above to re-normalise the data, excluding the effect of substrate folding.

First, this folding detection / moving variance procedure was performed on control data taken without added Cdc48 to obtain an estimate of the standard deviation of $\hat{\boldsymbol{\sigma}}^2$ expected under normal conditions. For the ubiquitin chain construct and a 1 s moving window, this standard deviation was $\sigma = 0.183$. The algorithm was then repeated on the experimental data taken in the presence of Cdc48, and each point with $\hat{\boldsymbol{\sigma}}^2 \geq 1.549$. ($3\sigma$ from the expected mean, $\hat{\boldsymbol{\sigma}}^2 = 1$) was marked as non-processive activity. Finally, contiguous regions of non-processive activity with duration less than 1 s were discarded.

Automated detection of non-processive activity was performed on all constant-distance data taken on the ubiquitin chain (polyubiquitinated MBP-ubi) construct.

*Processive extrusion & retrograde movement detection*

Processive extrusion appears as a gradual decrease in the measured contour length, over a period of 0.1 - 10 s on average. Retrograde movement appears as a complementary increase in the measured contour length, which may be sharp or similarly gradual. Each of these types of length changes may be evaluated using a series of Shewhart control charts constructed for each measured molecule.

Shewhart control charts provide a statistical framework for the quantification of ‘normal’ (‘in-control’) variability in one parameter describing a given process, and thereby enable detection of abnormal (‘out-of-control’) changes in that parameter[67]. In this way, provided an appropriate measurement parameter is selected, a single control chart may capture all the variability in possible extrusion and retrograde events above in a single set of statistical criteria.

In the context of the single-molecule experiments presented herein, the measurement parameter was chosen as the change in length $\Delta L$ at each time step $\Delta t$. For an ‘in-control’ process (*i.e.* a molecule tethered at constant force with no interacting partner), $\Delta$L is normally distributed about some mean $\mu$ and standard deviation $\sigma$. Note that this assumption of normality is not required for a Shewhart control chart, but does conveniently allow the same control limits to be used for detection of out-of-control events both above and below the process mean[68].

In practice, charts were constructed by segmentation of each length-time trace into regions of width $t$ = 0.2 s (≡ 50 data points at 500 Hz, the experimental measurement frequency after decimation). Each trace was segmented 10 times, where each set of segments was offset from the previous by 0.02 s. Overall, each molecule was thereby sampled at rate $f$ = 50 Hz ($\tau$ = 0.02 s) with sample size equivalent to $t$ = 0.2 s. This resampling procedure proved necessary to capture the wide variety of possible extrusion events. The high sample rate (above the Nyquist limit[69,70]) enables detection of the shortest, highest-frequency extrusion events. Equally, the large sample size enables detection of the longest extrusion, lower-frequency events, which are lost at lower sample sizes.

The same segmentation routine was performed for each molecule measured without added Cdc48 to obtain estimates of the parameters describing the ‘in-control’ process variability. For the conformationally-free construct, these were $\mu$ = - 0.11 nm and $\sigma$ = 5.180 nm.

Extrusion and retrograde events were then identified by comparison of each control chart with the following ’run conditions’. These conditions were adapted from those defined by Shewhart[67] and Western Electric[71] to differentiate between out-of-control regions with increasing mean (retrograde), and those with decreasing mean (extrusion). Extrusion events are those which fit any of the following criteria:

- 1 or more points beyond three standard deviations ($\Delta L < \mu - 3\sigma$);
- 2 of 3 consecutive points beyond three standard deviations ($\Delta L < \mu - 2\sigma$);
- 4 of 5 consecutive points beyond one standard deviation ($\Delta L < \mu - \sigma$);
- 7 or more consecutive points below the mean ($\Delta L < \mu$);

Retrograde events are defined using the same criteria above the mean ($\Delta L > \mu + 3\sigma$, *etc.*).

After run condition evaluation, the out-of-control regions identified in each chart were merged. As Shewhart charts are sampled at a lower rate than the raw data acquisition, the identified regions do not correspond exactly with the onset of Cdc48 activity in time. To optimise the start and end coordinates of each identified region, each pair of data points within 1 s (time) and 5 nm (length) of those initially defined obtained by Shewhart analysis were selected as candidates. A linear regression was fit to the data defined by each candidate start / end point. For extrusion events, the start/end pair with the best-fitting (highest $r$-value) regression were selected as the final start/end coordinates. Finally, extrusion events with duration less than 0.2 s, length less than 18 nm or velocity less than 5 nm $s^{-1}$; and retrograde events with duration less than 0.02 s or length less than 18 nm; were discarded.

In cases that a retrograde region overlapped an extrusion region, the extrusion region was split into two around the centre of the identified retrograde event. This enabled detection of repetitive extrusion-slip-extrusion sequences which would otherwise have been marked as one single event.

Unlike in the detection of non-processive activity, initial filtering of folding / unfolding events is unnecessary as the constructed Shewhart charts are not directly dependent on the variance on the measured length and therefore do not need to be re-normalised upon folding of the substrate.

For extrusion events (**Figure 2E**), velocities were computed as the slope of a linear regression of the length-time trace in a moving window with width $t$ = 1.5 s. As retrograde events are (on average) significantly quicker than extrusions, retrograde velocities (**Figure 3E**) were computed as the slope of a single linear regression over the entire retrograde event.

Automated detection of processive extrusion and retrograde movement events was performed on all constant-distance data taken on the 'conformationally-free' (polyubiquitinated MBP-ubi-ubi-MBP) construct.

*Survival analysis and rate calculations*

For both sets of automatically-analysed data (non-processive activity on the ubiquitin construct, and processive activity on the conformationally-free construct), a series of survival analyses were performed. Four time parameters were defined for each data set: the time between the beginning of the measurement and the first detected event, $t_{initiation}$; the time between the first and last detected events, $t_{total\ active}$; the duration of detected events, $t_{active}$; and the time between detected events, $t_{inactive}$. For each time coordinate, a survival function was defined using a Kaplan-Meier estimator[72]. For survival functions exhibiting exponential behaviour, the decay rate, $k$, for each survival function may be determined from an exponential fit to that function, with the form:

$$f(t) = Ae^{-kt},$$

where the scaling factor A = 1, since the fit is to a survival curve which represents a probability density function. In certain cases ($t_{inactive}$ in the processive data set $t_{active}$ and $t_{inactive}$ in the non-processive data set), the survival functions instead exhibit a biexponential distribution of the form

$$f(t) = A_1e^{-k_1t} + A_2e^{-k_2t},$$

where in this case $A_1 + A_2 = 1$.

To determine whether a mono- or bi-exponential model was optimal, the two models were compared for each survival curve fit by jack-knifing ('leave-one-out') cross-validation[73,74]. In brief: for each data set, each data point was removed one-by-one; the remaining $N - 1$ data points were fit with each model; and in each case, the likelihood that the $N$th data point had been left out of the set was computed. Multiplying the likelihoods obtained at each step then gives a robust measure of the goodness-of-fit of each model, which can be directly compared. The model with the greatest likelihood was selected.

Note that in cases were a bi-exponential model was optimal, we cannot say with confidence which of the two components corresponds to the theoretical rates given above. In these cases, the two component rate constants are simply labelled 1 and 2.

Fits for both non-processive and processive activities are shown in **Extended Data Fig. 4G** and **Extended Data Fig. 12E** respectively. Finally, a fifth survival function was defined incorporating both $t_{initiation}$ and the molecules in which no activity was detected. This is shown for non-processive activity in **Extended Data Fig. 4G**, and for processive activity in **Extended Data Fig. 12A**.
